# A computational neuroimaging account of impulsive premature decision-making

**DOI:** 10.64898/2026.01.18.699652

**Authors:** David M. Cole, Andreea O. Diaconescu, Christoph D. Mathys, Tina Wentz, Zoltan Nagy, Erich Seifritz, Boris B. Quednow, Lionel Rigoux

## Abstract

Impulsivity is a multidimensional construct with distinct implications for the pathophysiology and treatment of various neuropsychiatric conditions, including substance use disorders, behavioural addictions, attention deficit hyperactivity disorder, psychosis, and personality disorders. One proposed behavioural subtype, *premature responding impulsivity* (PRI), appears to be influenced by neuromodulatory pathways frequently implicated in these disorders, in particular dopamine neurotransmission, known for its role in contingency learning. Still, the neurobiological basis of PRI in humans remains insufficiently understood. Here, we theorize that PRI reflects the brain’s capacity to adapt to environmental uncertainty. To test this hypothesis, twenty-four healthy adults (mean age 22.6 years; 12 females) completed a novel decision-making task featuring alternating stable and volatile probabilistic cue contingencies while undergoing functional magnetic resonance imaging (fMRI). A hierarchical Bayesian model estimated PRI as an urgency-to-respond process, whose parameters were dynamically modulated by volatility. These model-derived indices correlated with established trait impulsivity measures, supporting their construct validity. Model-based fMRI analyses identified a distributed cortico-subcortical network including anterior insula, dorsal anterior cingulate cortex, striatum, and monoaminergic midbrain regions, whose activity tracked within-trial PRI estimates as they evolved over time. Connectivity analyses further showed that high volatility enhanced interactions between subnetworks typically associated with promoting or inhibiting impulsive action. Together, these results outline a neurocomputational account in which environmental uncertainty modulates PRI through interacting brain circuits, offering a principled framework for further probing the transdiagnostic role of impulsivity across neuropsychiatric and neuropsychopharmacological contexts.

## Introduction

Impulsivity can be construed as a normal facet of behaviour, affording an individual the ability to act optimally in a potentially urgent situation whilst minimising the time and neurocognitive processing resources required to consider and compare all possible choices (Dickman, 1990; Stevens and Stephens, 2010). A familiar example is online purchasing. During periods of rapid price fluctuations, such as seasonal sales, delaying a purchase in order to compare vendors may yield a better deal, albeit at the risk of stock depletion and thus potentially missing the actual lowest price. In contrast, when prices are predictable, prolonged comparison offers less benefit and quickly committing to one of the first acceptable options is often a more efficient choice in terms of minimizing the time and search effort invested. In general, the ability to rapidly process incoming information and to quickly implement an optimal choice based on selective sampling of this information thus confers clear benefits for humans and other organisms capable of making sophisticated decisions in fluctuating environments (Tickle et al., 2023). However, in behavioural contexts where acting with urgency confers no benefit, or is perhaps even counterproductive, an excess of impulsive behaviour can prove disadvantageous. Following our example above, you may regret buying an item if you find it at a better price soon after. In this specific situation, the relative benefits of deciding quickly versus waiting for more decision-relevant information can shift over time based on unknown variables (e.g., available stocks, seasonal offers). Beyond this illustrative example, the ability to perceive and adapt to such contextual changes is a fundamental property of decision-making and a key mechanism of adaptive behaviour.

Societally, inopportune impulsivity typically manifests as behaviours often regarded as problematic in human interactions and which, in extreme cases, can carry severe legal consequences for an individual, despite presumably arising from very similar neurobiological processes to those underlying comparatively beneficial, or prosocial, rapid decision-making (Raine and Yang, 2006; Dalley et al., 2011). Clinically, maladaptive impulsivity underlies behavioural abnormalities seen in multiple neuropsychiatric conditions including substance use disorders, behavioural addictions, attention-deficit/hyperactivity disorder (ADHD), psychosis, and personality disorders (Winstanley et al., 2006; Dalley et al., 2011; Moutoussis et al., 2011a; Soloff et al., 2014; Zorrilla and Koob, 2019). However, impulsivity is a heterogeneous, multidimensional construct that needs to be decomposed into distinct facets to capture its diverse manifestations and associations with clinical and biological variables (Gustavson et al., 2020; Strickland and Johnson, 2021; Wormington et al., 2026). Clinical research therefore depends on the reliable and valid operationalisation of these facets. One such specific subtype of impulsivity, referred to as ‘premature responding impulsivity’ (PRI), weaves a common thread through a wide range of psychiatric disorders (for a review, see Dalley and Robbins, 2017). For instance, PRI is at the core of the ‘jumping-to-conclusions’ bias of individuals with psychosis or schizophrenia, who may assign unwarranted significance to unexpected sensory events (Moutoussis et al., 2011b; Huddy et al., 2013). The importance of PRI is also evident in ADHD and in behavioural or substance addictions, where individuals show an impaired ability to withhold an action until enough information is available in unpredictable or dynamic settings (Winstanley et al., 2006; Voon et al., 2014; Banca et al., 2016; Morris et al., 2016; Dalley and Ersche, 2019).

On the biological level, PRI is conspicuous by the relative specificity of its neurochemical underpinnings (Dalley and Roiser, 2012). Indeed, unlike other major impulsivity subtypes such as motor impulsivity and delay discounting of reward, PRI largely appears to be modulated in opposing directions by the activity of dopaminergic and serotonergic pathways (Dalley and Roiser, 2012). More precisely, the literature suggests that increasing systemic dopamine (DA) levels tends to increase PRI behaviour, whereas increasing serotonin (5-HT) systemically attenuates PRI (for a review, see Dalley and Roiser, 2012). This dissociation parallels long-held theories that the effects of DA are most noticeable in contexts of decision-making and action execution relevant for acquiring reward, while those of 5-HT may be more prominent in contexts of behavioural inhibition and loss avoidance, although much nuance is required when translating this hypothesised dissociation to empirical observations (Daw et al., 2002; Fletcher et al., 2007; Rogers, 2011; Guitart-Masip et al., 2012, 2014; Seymour et al., 2012; den Ouden et al., 2013; Cohen et al., 2015; Moran et al., 2018; Rothenhoefer et al., 2019). The fact that both of these neuromodulatory systems are selectively targeted by medications commonly used to treat the symptoms of neuropsychiatric disorders in which PRI is thought to play a role reinforces the importance of those monoaminergic pathways for the expression of this form of impulsive behaviour.

Importantly, a growing volume of literature highlights the mechanistic influence, at various levels of the human brain’s decision-making hierarchy, of the same monoaminergic systems in mediating the neurobehavioural response to shifts in environmental uncertainty, or “volatility”, (Behrens et al., 2007a; Iglesias et al., 2013; Payzan-LeNestour et al., 2013; Marshall et al., 2016; Diaconescu et al., 2017). These overlapping contributions of DA and 5-HT suggest a tight link between impulsive decision-making and dynamic adaptation to uncertainty. While previous research started to explore this link, (Huettel et al., 2005; Franken et al., 2008; Tzagarakis et al., 2013; Voon et al., 2014; Tanovic et al., 2018), the precise role and neural basis of the influence of environment volatility on PRI remains surprisingly under-studied, despite its relevance to biological psychiatry.

Here, we aim to clarify this gap by proposing that individual differences in PRI are related to the behavioural and neural mechanisms by which we continuously adapt to the uncertainty in the statistical structure of our environment. To test this hypothesis, we apply a principled, model-based approach (Stephan et al., 2015) to characterise PRI and its neurophysiological correlates in humans. More precisely, we designed a novel paradigm evoking a dynamic decision-making process combining evidence accumulation and uncertainty belief updating. Building on a well-established, computational framework capturing learning of stable versus volatile contexts (Behrens et al., 2007b; Mathys, 2011; Mathys et al., 2014), we developed a computational model allowing us to formally test how PRI dynamics articulate with (Bayesian) statistical learning under uncertainty, thus capturing the ability to flexibly tune behavioural strategies to the volatility of the environment.

We hypothesised, firstly, that individuals expressing higher baseline levels of impulsive behaviour would display a reduced ability to adapt to shifts in uncertainty caused by alternating phases of relative stability and volatility of the learning environment. Secondly, we expected an overlap between the neural signatures – assessed using functional magnetic resonance imaging (fMRI) – of PRI behaviour and beliefs related to uncertainty tracking, particularly in dopaminergic circuits. Accordingly, we appraise our computational neuroimaging framework on three fronts: First, we establish construct validity by relating model-derived PRI indices to well-validated trait impulsivity scales. Second, we determine whether these indices capture the predicted impairments in adapting to shifts between stable and volatile contingencies. Third, we test whether the neural signatures of PRI map onto established reward- and motivation-related circuitry, and scale systematically with the statistical structure of the environment.

## Results

We invited 23 healthy volunteers to perform a new behavioural task designed to assess their impulsive responding in an environment with time-varying contingencies (see Fig. 1). We then fitted and compared a suite of computational models capturing various possible influences of the task statistical structure, implemented using a hierarchical Bayesian learning scheme, on response dynamics, implemented as an evidence accumulation process (for exhaustive methodological details see *Methods* and the Supplementary Materials).

**Figure 1.**
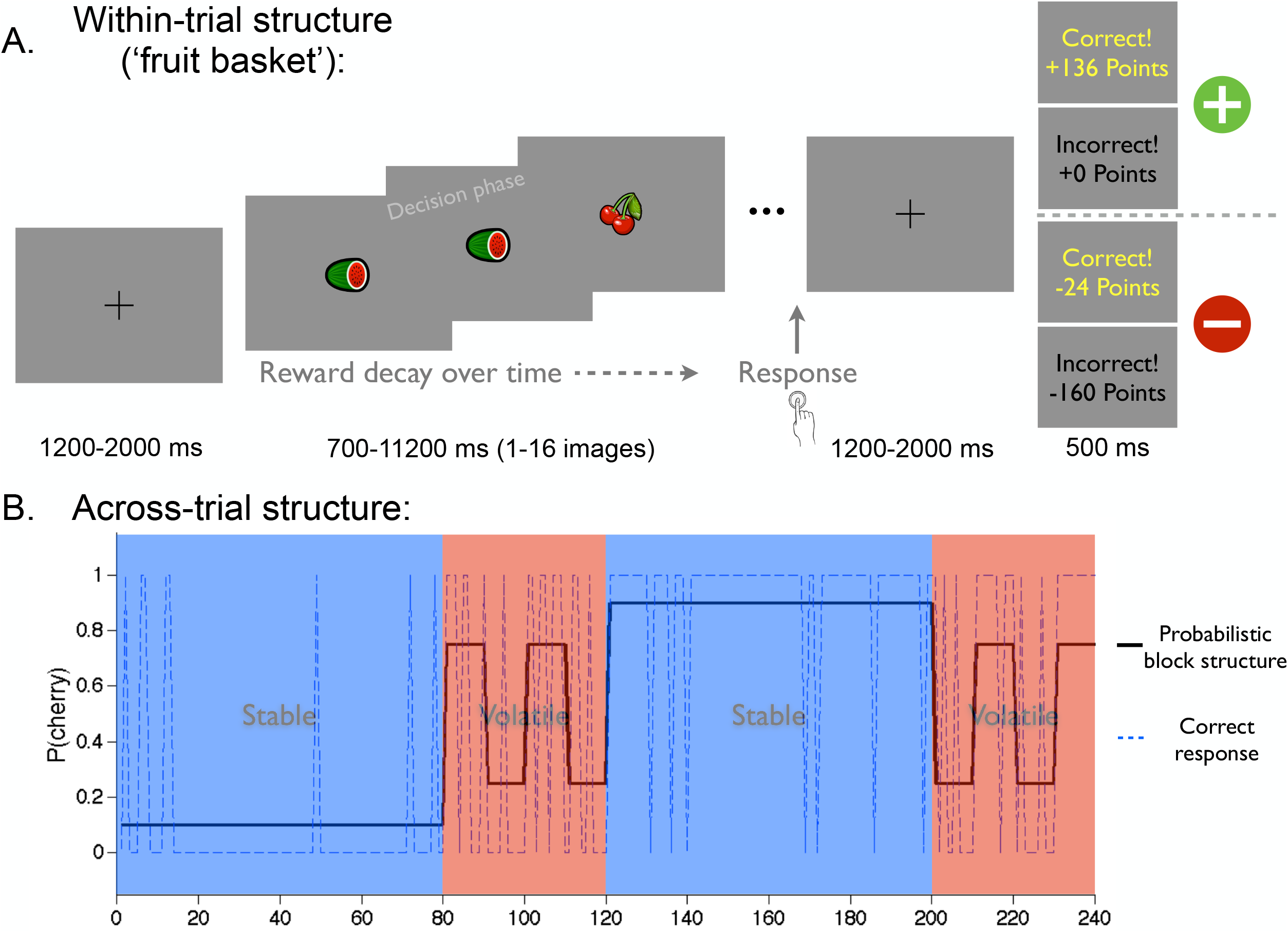
The time-varying uncertainty (TVU) task. (*A*) *Within-trial* (“*fruit-basket*”) *structure*. Each round presents up to 16 cartoon fruits (cherries or watermelon; 380 ms on, 320 ms off). Baskets come from **Vendor X** (10 cherries : 6 watermelon) or **Vendor Y** (10 watermelon : 6 cherries). Participants press with the right index (cherries) or middle finger (watermelon) as soon as they decide which fruit is more frequent (10 : 6 corresponds to p = 0.625 vs 0.375). The response ends the trial; after a jittered 1.2–2 s fixation, feedback (500 ms) shows accuracy and points. Reward decays linearly with each viewed cue: **Gain version** +160 to +100, **Loss version** -0 to –60; errors or time-outs (>320 ms after the last cue) yield +0 or –160 points. Inter-trial intervals are jittered 1.2–2 s. *(B) Across-trial structure*. Each session (240 Gain and 240 Loss trials) comprises four pseudo-blocks: 80 Stable, 40 Volatile, 80 Stable, 40 Volatile. In Stable blocks, one vendor predominates with p = 0.9/0.1; in Volatile blocks probabilities are 0.75/0.25 and the high-probability vendor flips every 10 trials (blue = Stable, red = Volatile). Early responding is optimal when the environment is Stable; waiting for additional cues is advantageous under Volatility, creating a reward-evidence trade-off thereby targeting premature responding impulsivity.

### Evidence accumulation dynamics reflect impulsive behaviour

In every trial, participants had to estimate which type of fruit is likely to appear more frequently in the sequence. At each cue presentation within the trial, they could decide to either indicate their choice – the sooner the larger the benefit – or wait and gather more information. This tendency to temporize, counterbalancing evidence accumulation, was formalized in our model by the parameters *w*_0_, reflecting the basal propensity to wait, and *u*, capturing a sense of urgency as the trial went by. Accordingly, *w*_0_ positively correlated with average choice latency (CL; r = –0.83, p < 0.001).

We also computed an Impulsivity score (*I-score*) and Efficiency score (*E-score*), data-driven metrics that respectively reflect impulsivity and overall response efficiency while controlling for potential speed-accuracy trade-offs. Accordingly, the *I-score* selectively negatively correlated with the tendency to wait (*w*_0_: r = –0.71, p < 0.001; *u*: r = 0.11, p = 0.629), while the *E-score* mapped preferentially with the time pressure parameter (*u*: r = –0.64, p < 0.001; *w*_0_: r = –0.42, p = 0.044), highlighting the capacity of the model to extract distinct characteristics of information sampling behaviour and thus discriminate the impulsive dimension of premature responding.

Altogether, those results consolidate the idea that *w*_0_ mirrors the PRI we wish to characterize. We therefore checked if this parameter also reflected psychometric measures of impulsivity based on self-report (see *Methods* – *Questionnaires*). While *w*_0_ did not significantly correlate with the total *Barratt Impulsiveness Scale* (BIS-11) score (r = – 0.33, p = 0.123), it however did so with the *attentional impulsivity* subscale (r = – 0.43, p = 0.041). The parameter *w*_0_ also correlated with *UPPS Impulsive Behavior Scale* scores (r = – 0.43, p = 0.041), further confirming that this computational trait is related to general constructs of impulsivity.

### Environment volatility modulates evidence accumulation behaviour

We designed our task such that the outcome of each trial (*i*.*e*., the fruit most likely to dominate the sequence of cues) could be, to some degree, predicted from the outcome of previous trials. However, the mean and variance of this trial-to-trial contingency was systematically varied to require participants to constantly learn, and to induce phases where learned predictions were more or less reliable.

To assess the influence of the environment predictability on impulsive behaviour, we first compared response latencies between stable and volatile phases of the task. When the contingencies were more stable, participants responded significantly faster (stable: CL = 9.3±3.3 cues; volatile: CL = 11.4±3.2; Δ CL = 2.1 cues, t(22) = 4.06, p < 0.001), while also making fewer errors (stable: errors = 14.6±4.3 %; volatile: errors = 19.0 ± 9.3 %; Δ errors = 4.3 %; t(22) = 2.51, p = 0.020), suggesting that participants leveraged the statistical structure of the task to adapt their decision-making.

One interpretation of this behavioural shift is that outcome predictions, based on past learning, are used to improve performances. Yet, we found no differences in E-scores between the two phases (stable: 1.47±1.14; volatile: 1.46±0.59; Δ E-score= – 0.01, t(22) = – 0.03, p = 0.977), suggesting no changes in task efficiency. Strikingly, the I-score significantly fluctuated during the task (stable: –1.01±1.29; volatile: –2.37±1.62; Δ I-score = –1.36, t(22) = –7.22, p < 0.001), pointing to a volatility-induced modulation of impulsive responding.

### Computational dissection of the behavioural adaptation to the task structure

In order to better understand those complex results, we designed a set of computational models implementing various hypotheses relating to the impact of learned outcome predictions and volatility estimates, based on a hierarchical Bayesian scheme, on the evidence accumulation dynamics (see *Methods – Behavioural modelling* and Table 1). Briefly, Bayesian model selection clearly identified *M*12: 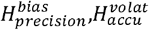 as the best explanation of the responses observed in our task (Ef = 0.55, *pxp* = 1). This model identifies two critical mechanisms.

**Table 1.** Model space considering two factors influencing canonical behaviour during the task. Asterisk (*) denotes winning model identified via group Bayesian Model Selection.

|  |  | Hypothesis iterations based on which decision model parameter is predicted to be influenced by log-volatility |  |  |  |  |
| --- | --- | --- | --- | --- | --- | --- |
| | | $H_{none}^{volat}$ | $H_{u0}^{volat}$ | $H_{ut}^{volat}$ | $H_{accu}^{volat}$ | $H_{bias}^{volat}$ |
| Hypothesis iterations based on predicted influence of HGF prior on evidence accumulation | $H_{none}^{bias}$ | $M_1$ | $M_2$ | $M_3$ | $M_4$ | -- |
| | $H_{fixed}^{bias}$ | $M_5$ | $M_6$ | $M_7$ | $M_8$ | $M_{13}$ |
| | $H_{precision}^{bias}$ | $M_9$ | $M_{10}$ | $M_{11}$ | $M_{12}^*$ | $M_{14}$ |

First, the current belief about the most likely outcome serves as a starting point of the information sampling process in a given trial. More precisely, we found that the mean outcome prediction biased the evidence accumulation (*β*_*µ*_ = 2.96±1.7; t(22) = 8.51, p < 0.001), but this influence was reduced when the belief variance was high (*β*_*v*_ = – 155.7±106.6; t(22) = – 7.00, p < 0.001). In other words, the contingencies learned from past trials are used as (precision-weighted) priors to inform future decisions. This initial bias contributed to the increase in performance observed in the stable phase, as a stronger bias (*β*_*µ*_) was associated with larger improvement in choice latencies (r = – 0.55, p = 0.006), error rates (r = – 0.70, p < 0.001), and E-scores (r = 0.81, p < 0.001) but not impulsivity-related I-scores (r = –0.02, p = 0.944).

The second mechanism identified by our model selection is a modulation by volatility of the information sampling dynamics. More specifically, we found that participants accumulated evidence faster when the volatility was high (*ξ*_*α*_ = 3.9±4.6, t(22) = 4.11, p < 0.001). The strength of this modulation was reflected in the task phase-induced differences in errors (correlation *ξ*_*α*_ × Δ error %: r = 0.71, p < 0.001) and efficiency (correlation *ξ*_*α*_ × Δ E-score: r = – 0.51, p < 0.013) but not in choice latency (*ξ*_*α*_ × ΔCL: r = 0.21, p = 0.333), which surprisingly did not translate significantly to an increase in impulsivity (*ξ*_*α*_ × Δ I-score: r = 0.25, p = 0.244). In summary, participants relied more on environmental cues in volatile phases, an effect opposing and complementing the effect on learned priors in stable phases.

Interestingly, the amplitude of the aforementioned reliance on learned expectations in the stable phase was strongly correlated with this volatility-induced modulation of sampling (r = 0.67, p < 0.001), suggesting that those effects are two sides of the same coin. One interpretation is that these complementary dynamics reflect a long-term strategy: by saving time in the volatile phase, one can allocate more resources to the more profitable stable phase of the task. This interpretation is supported by the fact that both effects were associated with higher reward rates (correlations: *β*_*µ*_ × points/cue: r = 0.64, p < 0.001; *ξ*_*α*_ × points/cue: r = 0.48, p < 0.02) and lower delay discounting (correlations: *β*_*µ*_ × KirbyK: r = – 0.54, p = 0.007; *ξ*_*α*_ × KirbyK: r = – 0.67, p < 0.001).

Altogether, these results highlight the highly adaptive nature of response impulsivity. More precisely, our computational approach identifies a multifaceted modulation of the evidence sampling strategy by (Bayesian) beliefs about the statistical structure of the environment.

Finally, we explored the relation between trait impulsivity reflecting impulsive behaviour (especially PRI) in daily life, as captured by *w*_0_, and the amplitude of the dynamic modulation of PRI by the environment we identified above. We found that higher basal impulsivity was associated with a reduced volatility influence on decisions (correlations: *w*_0_ × *β*_*µ*_ : r = – 0.72, p < 0.001; *w*_0_ × *ξ*_*α*_ : r = – 0.57, p < 0.001), suggesting that participants with higher trait impulsivity also failed to adapt their impulsive responding to the changes of the environment.

### Neural correlates of impulsive decision-making

To better understand the neural basis of PRI and its modulation by volatility, we adopted a model-based analysis of the fMRI data acquired during the task (see *Methods – fMRI analysis*).

A negative contrast on the cue-by-cue dynamic waiting signal, *w*^*t,k*^ – reflecting more impulsive decision-making – revealed significant, mostly bilateral activations across multiple brain regions, notably encompassing the anterior insula, the dorsal anterior cingulate cortex, the dorsolateral prefrontal and superior frontal cortices, the precuneus, the caudate, the midbrain, and the subthalamic nucleus (see Fig. 2A and Table 2 for exhaustive results).

**Table 2.** Brain regions significantly encoding waiting impulsivity (1/*w*). AI = anterior insula; dlPFC = dorsolateral prefrontal cortex; dACC = dorsal anterior cingulate cortex; STN = subthalamic nucleus; HPC = hippocampus; STS = superior temporal sulcus; L = left hemisphere; R = right hemisphere.

| Brain region | Cluster size<br>(2 mm<sup>3</sup> voxels) | p (whole brain<br>FWE-corrected) | t-value of peak<br>voxel | x, y, z co-ordinates<br>(MNI space) |
| --- | --- | --- | --- | --- |
| R AI extending to dlPFC | 2900 | < 0.001 | 15.94 | 40, 22, 2 |
| Bilateral lingual gyri extending to occipitotemporal and inferior parietal cortices and dorsal cerebellum | 15176 | < 0.001 | 14.80 | -8, -78, -2 |
| Bilateral dACC | 788 | < 0.001 | 14.15 | 4, 16, 48 |
| L AI | 606 | < 0.001 | 12.41 | -34, 26, -2 |
| Bilateral midbrain ext. to STN, thalamus, caudate and R HPC | 1563 | < 0.001 | 10.68 | 10, -24, -12 |
| R superior frontal cortex | 166 | < 0.001 | 10.25 | 36, -4, 60 |
| R STS | 124 | < 0.001 | 9.60 | 50, -22, -8 |
| L dlPFC | 178 | < 0.001 | 9.24 | -28, -4, 44 |
| L superior frontal cortex | 314 | < 0.001 | 9.17 | -28, -8, 56 |
| R precuneus | 47 | < 0.001 | 8.10 | 10, -42, 52 |
| L posterior thalamus/HPC | 45 | < 0.001 | 7.85 | -20, -28, -2 |

**Figure 2.**
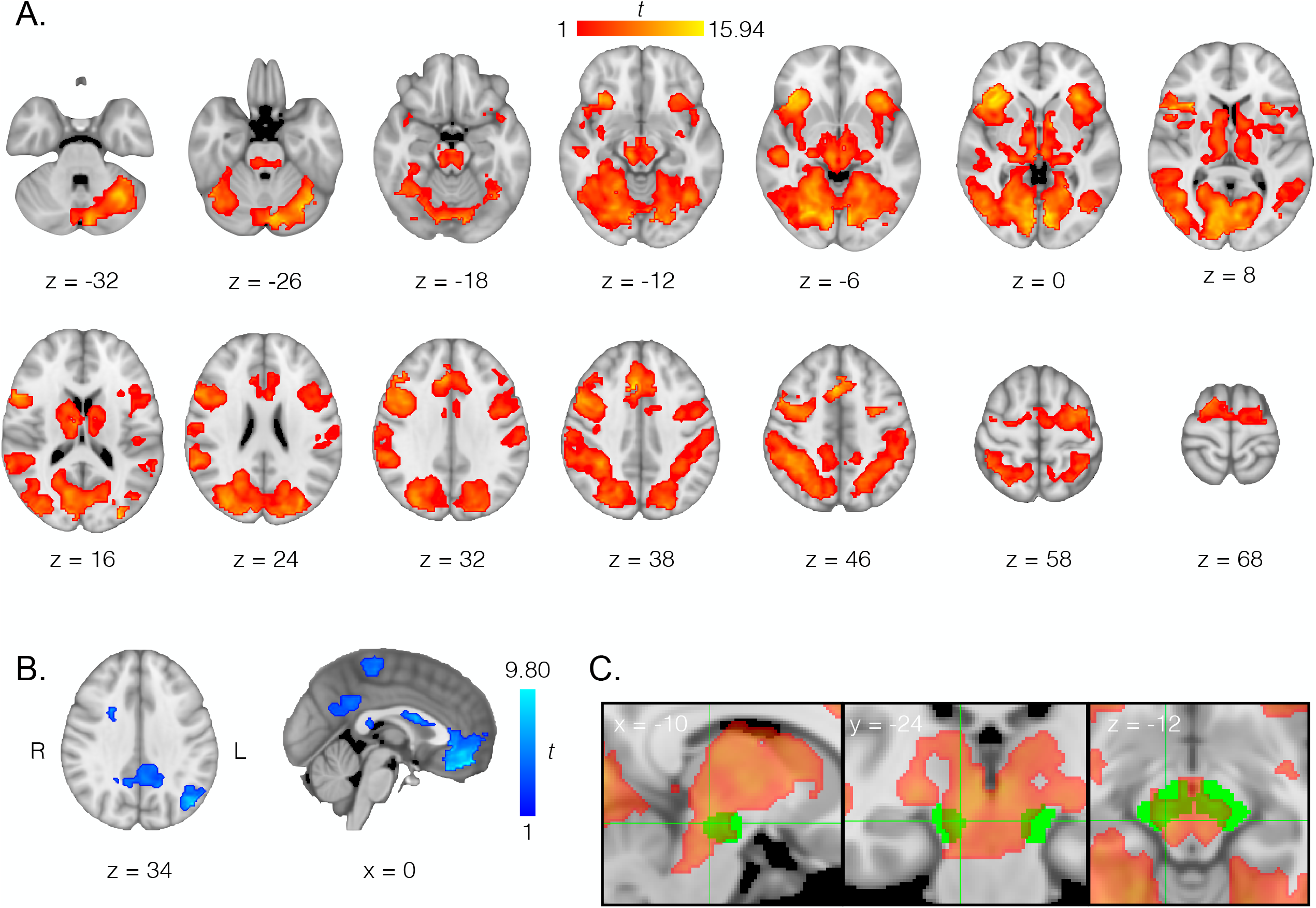
Network of brain regions related to impulsive behaviour during the task. (A) Brain regions where BOLD positively correlated with PRI behaviour; that is, negatively correlated with waiting behaviour *w* (whole-brain cluster-level FWE-corrected p < 0.05, cluster extent threshold *k* = 92 voxels). (B) Brain regions where BOLD *negatively* correlated with PRI behaviour, that is positively correlated with waiting behaviour *w* (whole-brain cluster-level FWE-corrected p < 0.05, *k* = 92). R = right, L = left. NB: axial slices are displayed in line with radiological orientation conventions (L/R hemisphere flipped in figure). Colour bars represent *t*-statistic effect sizes shown in brain image overlays. Statistical maps overlaid on background image of a T1 template in MNI152 standard space. (C) Additional overlay of an *a priori* mask of the SN/VTA (green), visibly overlapping with the activation map from ‘A’ above (red-yellow, reduced opacity); crosshairs denote peak midbrain activation. For detailed visualisation purposes the overlaid statistical images and background anatomical image in all panels are presented in 1mm^3^, rather than 2mm^3^, isotropic resolution.

When considering, conversely, the effects of a *positive* contrast on *w*, meaning increased activation when decision-making is *less* impulsive, we found significant activations in a different set of brain regions, notably including the bilateral ventromedial prefrontal cortex, the posterior cingulate cortex, and the precentral gyri (Fig 2B; see Table 3 for details).

**Table 3.** Brain regions significantly *negatively* encoding waiting impulsivity (1/*w*), or positively encoding waiting behaviour (*w*). AI = anterior insula; vmPFC = ventromedial prefrontal cortex; mOFC = medial orbitofrontal cortex; PCC = posterior cingulate cortex; ITS = inferior temporal sulcus; L = left hemisphere; R = right hemisphere; ^§^denotes two clusters where significance is maintained when using a more conservative, whole-brain voxel-level FWE-correction (p < 0.05, extent threshold 45 voxels or greater, cluster size in italics, data in other columns remain.

| Brain region | Cluster size<br>(2 mm<sup>3</sup> voxels) | P (cluster-level<br>FWE-corrected) | t-value of peak<br>voxel | x, y, z co-ordinates<br>(MNI space) |
| --- | --- | --- | --- | --- |
| Bilateral vmPFC and mOFC<sup>s</sup> | 1160; *412* | < 0.001 | 9.80 | -2, 44, -18 |
| L inferior parietal/occipitoparietal cortex<sup>s</sup> | 554; *125* | < 0.001 | 8.14 | -44, -70, 36 |
| Bilateral dorsal caudate extending into white matter | 1196 | < 0.001 | 7.41 | -2, 6, 18 |
| R dorsal posterior cerebellum | 128 | 0.014 | 7.11 | 34, -82, -34 |
| R hippocampus ext. to temporo-occipital fusiform/lingual gyri | 189 | 0.002 | 6.82 | 38, -42, -2 |
| Bilateral precentral and postcentral gyri/somatomotor cortex | 712 | < 0.001 | 6.73 | -6, -28, 60 |
| Bilateral PCC/precuneus | 803 | < 0.001 | 5.78 | 10, -54, 32 |
| L ITS | 192 | 0.002 | 5.64 | -62, -12, -16 |

Following up on the neuromodulatory hypothesis of PRI, we refined our analysis utilizing anatomically defined masks of the substantia nigra/ventral tegmental area (SN/VTA; Bunzeck and Düzel, 2006), the dorsal raphe nucleus (Beliveau et al., 2015), and the locus coeruleus (Keren et al., 2009) to try and detect blood-oxygenation level-dependent (BOLD) markers of dopaminergic, serotoninergic, and noradrenergic influences in the expression of impulsive behaviour. While significant activation was detectable in the dorsal raphe (p < 0.05, cluster-level FWE-corrected for small volume), its overlapping extent with the mask was marginal (peak *t* = 6.30; p = 0.005; x = 4, y = -30, z = -8; 8% overlap with the mask volume), and no cluster was present in the locus coeruleus. However, as already evident in the whole brain results, we found significant representations of waiting impulsivity in the SN/VTA (peak *t* = 10.23; p < 0.001; x = 10, y = -24, z = -12; 43% mask volume overlap; Fig. 2C). Altogether, these targeted tests highlight the potential relevance and specificity of dopaminergic activity in the regulation of PRI.

### Neural basis of context-dependent regulation of impulsivity

Our computational analysis revealed that participants learned the task structure, allowing them to guess faster the most likely outcome of a trial and therefore improve their performance. Replicating previous studies, we found that this (hierarchical) learning process engaged striatal regions, critical for the encoding of (precision-weighted) predictions errors (Iglesias et al., 2013; see also Supplementary Materials – Fig. S1 and Tables S1-2). In our model, the belief resulting from this learning process yields a decision bias, *λ*^*t*^, which allows the agent to bootstrap their evidence accumulation strategy. Accordingly, a follow-up fMRI analysis shows that this decision bias downregulated the activity of various regions related to impulsive responding, including the right anterior insula (whole brain level) and the midbrain SN/VTA (small volume correction; see Supplementary Fig. S2 and Table S3 for details). Further, belief precision, which weighted both prediction errors and the decision bias, was encoded in regions also negatively encoding PRI (Fig. S3). In other words, those brain regions where BOLD activation increased with PRI also disengaged when prior information could be relied upon; a modulation possibly mediated by dopamine-dependent learning of the task structure.

### Volatility-dependent connectivity of the PRI network

Another behavioural mechanism we identified with our model is a boosting influence of environmental uncertainty on the evidence accumulation dynamics. In order to probe more directly the neurobiological basis of this relationship, we conducted a psychophysiological interaction (PPI) analysis (see *Methods – PPI analysis*) to assess the influence of the task volatility (*i*.*e*., stable vs. volatile blocks) on the functional connectivity of the right anterior insula which, as shown above, preferentially encoded decision impulsivity.

This approach identified numerous regions that showed higher functional integration with the anterior insula in volatile phases, including the posterior cingulate cortex, the precuneus, the ventromedial prefrontal cortex, the medial orbitofrontal cortex, and pre- and post-central gyrus regions within the somatomotor cortex (Fig. 3; Supplementary Table S4). Interestingly, this functional network partially overlapped with the brain regions negatively encoding PRI (Fig. 2B and Table 4), supporting the importance of this competing network in moderating the expression of decision impulsivity at the neural level.

**Figure 3.**
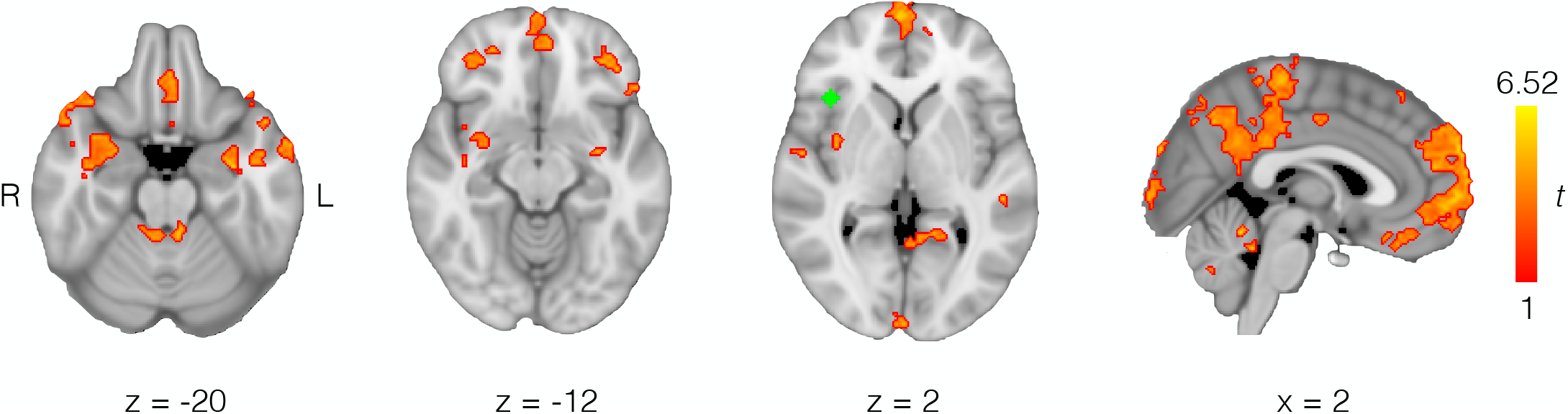
Brain regions displaying a volatility-dependent psychophysiological interaction effect with the insula-centred PRI network (see Fig. 3). Those regions increased their functional connectivity with the (pro-impulsive) right anterior insula (5 mm radius seed mask shown in green) during the volatile periods of the task (whole-brain cluster-level FWE-corrected p < 0.05, extent threshold *k* = 66). Colour bar represents t-statistic effect sizes shown in brain image overlay. R = right, L = left. NB: axial slices are displayed in line with radiological orientation conventions (L/R hemisphere flipped in figure). Binary and statistical maps overlaid on background image of a T1 template in MNI152 standard space.

## Discussion

Using a novel task requiring both evidence aggregation and contingency learning, we explored the influence of environment predictability (or volatility) on premature response impulsivity (PRI) – a facet of impulsive decision-making. Combining computational and fMRI approaches, we showed that the task structure has a dynamic, multifactorial influence on PRI, which is supported by a complex cortico-subcortical network that is adaptively modulated by the environment predictability. The activation of SN/VTA and caudate regions related to both contingency learning (precision-weighted prediction errors) and decision-making impulsivity (*w*) further substantiate the hypothesis that dopamine plays a critical role in mediating the adaptive nature of PRI.

Aberrant decision-making behaviours, of which impulsivity constitutes a key example, are of central interest to diverse branches of neuroscience, such as those dedicated to the study of psychiatric, neuroeconomic and neuroethical concepts and phenomena (Moeller et al., 2001; Glimcher et al., 2007; Henry and Plemmons, 2012; Lee, 2013). Despite this broad relevance, approaches to characterising the behavioural processes and neurophysiological mechanisms underlying impulsivity – as with the vast majority of symptom patterns targeted by clinical neuropsychiatric research questions – have as yet largely failed to advance beyond the descriptive towards providing validated, principled accounts of these phenomena (Stephan et al., 2015; Iglesias et al., 2017). In this study, we therefore aimed to mechanistically model such behaviours and characterise them in relation to more established cognitive-behavioural constructs and concepts. Our research focussed on PRI as a key subtype of impulsive behaviour, which to date has largely been investigated using animal models (e.g., Robbins, 2002; Fletcher et al., 2007; Dalley and Roiser, 2012). To this end, we applied a novel combination of model-based behavioural and fMRI techniques in order to illustrate the functional neurobiology of PRI in decision-making in human participants.

### Modelling PRI under uncertainty

Importantly – and contrasting with the findings of a small collection of prior computational studies exploring the mechanistic relationship between impulsivity and uncertainty (Averbeck et al., 2013; Paliwal et al., 2014, 2019) – we did not specifically predict that increased PRI would be associated with higher estimates of environmental uncertainty. Contrary to these studies, our paradigm was designed to reward rapid decision-making behaviour in stable environments and more deliberative behaviour in volatile environments. Moreover, the computational contexts in which these studies examined the relationship between impulsivity and uncertainty was either very different to our current framework (Averbeck et al., 2013), or did not involve consideration of PRI specifically (Paliwal et al., 2014, 2019). Therefore, we instead initially predicted that higher trait-level PRI would reduce the impact of volatility on decision-making behaviour.

Building on established approaches modelling hierarchical probabilistic decision-making under uncertainty (Mathys, 2011; Iglesias et al., 2013), we were able to successfully capture baseline trait-like and dynamic state-level impulsivity, then relate them respectively to behavioural variations, both across participants and decision-making task contexts. More precisely, we identified: (i) a general (trait) tendency to wait, related to self-report metrics of impulsivity; and (ii) a dynamic (state-dependent) strategy balancing the relative weight of prior beliefs *vs* visual cues, depending on the higher order statistics of the task contingencies (expectation, variance, and volatility of the environment). This formalism could therefore help with disentangling organic and context-dependent causes of impulsivity.

While our modelling approach relied on a normative, Bayesian implementation of the learning dynamics, the evidence accumulation and decision process was formalised in a more descriptive fashion. This allowed us to test various mechanistic hypotheses about how those two modules could interact based on empirical observations, rather than theoretical constraints or preconceptions. Based on our results, future work could incorporate theory-driven models of perceptual inference (e.g., FitzGerald et al., 2015; Denève and Jardri, 2016) to develop an integrated, normative model of impulsive responding prescribing how decision-making should optimally adjust to the structure of the environment.

### Model-based PRI in the context of established behaviour/personality metrics

Indeed, ‘impulsivity’, as commonly discussed, is a multifaceted construct, comprising a number of behavioural and personality subtypes and styles (Dickman, 1990; Evenden, 1999; Dalley and Robbins, 2017), which have partially differential clinical consequences (Strickland and Johnson, 2021; Wormington et al., 2026). Prior efforts to empirically compartmentalise and characterise the PRI subtypes have largely centred around animal research models, with a small amount of supporting human neuropharmacology and neuroimaging literature (Dalley and Roiser, 2012; Dalley and Robbins, 2017). Paradigms probing other primary subtypes of impulsive behaviour, namely motor impulsivity and temporal delay discounting of reward, have been relatively straightforward to translate to human neuropsychiatric research (Winstanley et al., 2006; Pine et al., 2010; Whelan et al., 2012; Vanderveldt et al., 2016). However, it is difficult to claim the same for PRI, which arguably describes a cognitively higher-order range of behaviours in humans than those employed in rodent models (Dalley and Roiser, 2012; Dalley and Robbins, 2017). Attempts have been made to adapt tasks such as the ‘5-choice serial reaction time task’ (5-CSRTT), which has been widely used to probe rodent models of PRI (see, e.g., Robbins, 2002), in order to test similar behaviour in humans (Voon et al., 2014, 2015; Worbe et al., 2014; Morris et al., 2016). While these studies support the translational validity of the 5-CSRTT across species in characterising premature responding behaviours (Voon et al., 2014; Worbe et al., 2014), which are known to be predictive of compulsive drug-seeking and other addictive behaviours in rodents (Belin et al., 2008; Voon and Dalley, 2016), much work is still needed to fully map the impulsivity subtypes across species (see e.g. Voon and Dalley, 2016). This is particularly true when higher cognitive domains are involved and when attempting to tackle neuropsychiatric conditions and their neurochemical substrates (Clark et al., 2006; Crockett et al., 2012; Voon et al., 2015; Banca et al., 2016; Tanovic et al., 2018; Kluwe-Schiavon et al., 2020; Zhukovsky et al., 2020). The task and associated computational modelling framework introduced in the current work provides a novel platform to dissect the mechanistic characterisation and ecologically valid classification of PRI symptomatology in humans. Indeed, our computational behavioural and neuroimaging approach is sensitive to individual differences in impulsivity, especially PRI, and provides a bridge to other established behavioural and personality questionnaire measures of impulsivity, such as (subscales of) the Barratt Impulsiveness Scale (BIS-11). However, our results also demonstrate a highly dynamic, context-dependent facet of impulsivity, prompting the need to clarify the distinction between mechanistic, behavioural, and neurobiological subtypes of impulsive behaviour beyond previous attempts (Dalley and Roiser, 2012), while at the same time underlining the suitability of a unified modelling approach for measuring and characterising PRI in humans.

### Neural signature of PRI

The results from our model-based fMRI approach, revealing an extended network of regions correlating with PRI behaviour, shows a high concordance with known cortical and subcortical brain systems previously associated with impulsivity, including fronto-insular and anterior cingulate cortex, as well as subthalamic and midbrain nuclei including the dopaminergic SN/VTA (Dalley and Robbins, 2017; Dalley and Ersche, 2018). Moreover, despite a sample size that might be regarded as small relative to many other contemporaneous neuroimaging investigations, we have found these effects to be extremely robust, in terms of their consistency across task reward contexts, their statistical effect sizes, and their neural spatial extent. Thus, our findings demonstrate the pertinence of our novel behavioural and computational paradigm to explore the neural basis of impulsivity.

The strong PRI-related activations in the anterior insula mirror the known critical role of this region in impulsive behaviours and interoceptive processing relevant for substance craving and addiction (Naqvi et al., 2007; Garavan, 2010; Churchwell and Yurgelun-Todd, 2013). In addition, our results corroborate previous findings implicating the subthalamic nucleus, not just in motoric aspects of “waiting” impulsivity (Morris et al., 2016; Mechelmans et al., 2017), but also specifically in “decisional” PRI, wherein electrical stimulation of this region was associated with increased PRI in terms of attenuated evidence accumulation in the beads task in individuals with obsessive compulsive disorder (OCD), thus ‘normalising’ their behaviour to be closer to that of healthy controls (Voon et al., 2017).

We also found activations related to *reduced* PRI in multiple regions including ventromedial prefrontal and posterior cingulate cortices – areas commonly identified as hubs of the brain’s “default mode network” (Raichle, 2015), which has been previously associated, negatively, with trait impulsivity (Li et al., 2013). Moreover, this brain network – and the posterior cingulate cortex in particular – is thought to be involved in internally-directed cognition and the ability to resist cravings (Bourque et al., 2013). Critically, we identified an almost identical set of regions that modulated their functional integration with the anterior insula as a function of the environment volatility. In line with the proposed role of the posterior cingulate cortex in the balance between internal vs external attention (Leech and Sharp, 2014), this network might be critical to dynamically and adaptively tune impulsive behaviour to adjust for fluctuations in beliefs and contextual changes.

Finally, we identified a tight link between contingency learning and evidence accumulation. Interestingly, the same SN/VTA-striatal circuit supporting prediction error encoding also correlated with computational trajectories of waiting impulsivity, suggesting a critical role of the basal ganglia (BG), and most likely dopamine, in mediating the ability to account for prior information during decision-making, which translated to an increased behavioural expression of impulsivity. Interestingly, as prediction errors (and therefore BG activity) were reduced when the task contingencies were more predictable, our results indirectly predict that impulsivity could be reflected by a relative decrease in BG function. This observation aligns, although with a different but not necessarily opposing interpretation, with an extensive literature arguing for a role of the BG in “braking” impulsive responding (e.g., Frank et al., 2007; Wessel and Anderson, 2023).

In summary, our findings emphasise a novel link between PRI behaviour and uncertainty, or volatility, in the decision-making environment. We have discovered that two functionally distinct networks – one supporting impulsive behaviour and centred on the anterior insula, anterior cingulate, midbrain and other regions, and one counteracting impulsive behaviour and centred on the posterior cingulate and ventromedial prefrontal cortex – become more functionally connected during periods of high volatility. Moreover, the network associated with impulsivity included brain regions critical for contingency learning, highlighting the tight integration of these behavioural mechanisms also at the neural level.

Altogether, our study provides computational and neurobiological evidence of a mechanistic link between impulsive behaviour and decisional uncertainty, calling attention to the adaptive, highly dynamic nature of impulsivity. Still, more work is needed to disentangle the causal relations and neural implementation of those inter-dependent (and thus statistically correlated) mechanisms.

### PRI and computational psychiatry

As was outlined in introduction, premature responding is known to be characteristic of the symptomatology of a range of neuropsychiatric conditions, many of which are routinely treated with drugs targeting monoaminergic systems. Studies with the Beads Task have shown, for example, that a ‘jumping to conclusions’ bias, hampering efficient evidence accumulation, exists in schizophrenia (Moutoussis et al., 2011b; Evans et al., 2015; Jardri and Denève, 2017). Interestingly, schizophrenia and paranoia seem to be robustly associated with belief updating disturbances and an overestimated volatility of the environment (Cole et al., 2020; Sheffield et al., 2022). According to our results, this elevated volatility prior, through its influence on evidence accumulation, might directly contribute to impulsive decision-making in those clinical conditions.

While first-episode psychosis is also associated with impulsive responding, this might be dependent on concomitant cannabis use (Huddy et al., 2013). Other studies investigating PRI in various types of substance use disorder have also found impairments across diverse PRI paradigms (Clark et al., 2006; Quednow et al., 2007a; Voon et al., 2014; Banca et al., 2016; Morris et al., 2016), while additional PRI aberrancies in evidence accumulation processes have been characterised in OCD (Banca et al., 2015; Voon et al., 2017) and posited in ADHD (Winstanley et al., 2006). The observation that PRI is modulated by the DA and noradrenaline reuptake inhibitor methylphenidate, which is often used to treat symptoms of ADHD, further supports its relevance in this case (Voon et al., 2015). Conversely, the above indications for PRI holding relevance for OCD symptomatology, as well as the finding that depleting the 5-HT precursor tryptophan in healthy individuals attenuates immediate loss aversion and promotes longer-term objective decision-making in a similar manner to sub-clinical depressive symptoms (Crockett et al., 2012), further promotes the neurochemically pervasive elements and thus the potentially ‘transdiagnostic’ utility of PRI measures in computational psychiatry (Dalley and Robbins, 2017; Iglesias et al., 2017). Despite these cumulative observations of its relevance, only a minority of previous investigations into PRI has sought to mechanistically model the behaviour being measured and, until now, no studies have deliberately harnessed the modulating role of environmental uncertainty in this effort. We thereby provide a model-based characterisation of the functional neurobiology of PRI for the first time in humans. This framework can thus pave the way for a raft of computational neuropsychopharmacological and clinical studies that will test hypotheses of the transdiagnostic potential of PRI directly.

Future experimental work should develop these initial proofs of concept further by investigating changes in PRI behaviour and neural computations elicited by targeted neurochemical modulations, as well as the influence of genetics and the potential identification of ‘neurocognitive endophenotypic’ markers of neuropsychiatric disorders (Robbins et al., 2012). For example, reuptake inhibitor pharmaceutical administration or amino acid depletion dietary protocols could be used, respectively, to increase or decrease the extracellular concentrations of DA or 5-HT (or noradrenaline), while the neurobehavioural effects of such modulations may interact with individual differences in the expression of genetic polymorphisms influencing these monoamine systems (Walderhaug et al., 2007; Forbes et al., 2009; den Ouden et al., 2013; Hennig et al., 2020). Ultimately, the incorporation of PRI behavioural and neuronal characterisation and classification techniques has the potential to further enhance the growing contribution of translational neuromodeling and computational psychiatric approaches to individualized medicine.

## Methods

### Participants

We collected data from 24 healthy volunteers (mean age = 22.6 years ± 2.2 s.d.; 12 female), who completed our behavioural paradigm while undergoing BOLD fMRI. Participants were recruited from the volunteer pool of the University Registration Centre for Study Participants, Department of Economics, University of Zurich, Switzerland. All participants were right-handed, had no history of neurological or psychiatric illness including brain injury or substance use disorder and all were non-smokers. Participants provided signed, informed consent and all study procedures were conducted concordantly with the standards of the Declaration of Helsinki. The study protocol was approved by the Ethics Committee of the Canton of Zurich (KEK-ZH 2010-0327). One participant (female, 20 years) experienced some dizziness while in the scanner and pressed the alarm button, thus terminating the experiment before the completion of data acquisition and precluding the inclusion of their behavioural and fMRI data in the described analyses.

### Behavioural paradigm

In the probabilistic, binary decision-making task (Fig. 1), each trial consisted of a sequence of 16 visual cues representing either a pair of cherries or a watermelon. One type of fruit was always dominating the sequence (10 vs. 6 cues) and participants were asked to select, using button presses, which fruit was the most frequent. Correct responses were either rewarded (‘Gain’ condition) or allowed to avoid punishments (‘Loss’ condition). Moreover, faster responses yielded higher rewards (or lower losses), forcing participants to decide, at each cue presentation, to either wait longer, and gather more information about the correct decision, or respond sooner, and maximise reward (respectively minimise loss) if correct. After indicating their choice, the cue sequence was immediately terminated and a feedback screen indicating the outcome was presented. Critically, the progression prescribing the correct (*i*.*e*., more frequent) fruit followed a pre-determined order (Fig. 1B, dashed line), similar to previously described ‘reversal learning’ tasks (Cools et al., 2002; den Ouden et al., 2013; Costa et al., 2015; Izquierdo et al., 2017) wherein contingencies varied at differing frequencies over time. In other words, the correct outcome at a given trial could be predicted from past outcomes, and the strength of this prediction fluctuated across task, being more reliable in ‘stable’ (80 trials) vs. ‘volatile’ (40 trials) pseudo-blocks (Fig. 1B, shaded areas).

Gain and loss conditions (240 trials each) were acquired independently. However, as no differences were found between those two framings, all parameters and data-driven measures were averaged across the two conditions (Table S5).

To ensure constant statistical power and maximize model identifiability (see below), a single, consistent contingency structure was selected (based on simulation) and programmed identically for both task runs across all participants. Order of completion of the Gain and Loss task runs and visual stimuli (‘cherries’ and ‘watermelon’) were counterbalanced across and within participants. The task was programmed with the Psychophysics Toolbox Version 3 (http://psychtoolbox.org) for Matlab (Natick, MA: The Mathworks Inc.).

### Behavioural modelling

#### Basic framework

Each trial was composed of a sequence of 16 cues representing either cherries or a watermelon, which were arbitrarily encoded as 0 or 1. The trial “type”, defined by the most frequent fruit in the sequence, and the participant response, were encoded accordingly. Finally, the on-screen feedback indicating whether the choice regarding trial type was correct or not was also coded as a binary outcome (*fbk* = 1 if correct, 0 otherwise). Choice latency (CL) denoted the index of the cue after which the subject provided their response, from 1 (right after the beginning of the sequence) to 16 (once the sequence is over and all cue stimuli *c*^1^-*c*^16^ have been presented on screen). Here, a complete lack of response is coded as a ‘null’ CL. The distribution of trial types across the experiment followed a hidden, but consistent, structure composed of alternating periods (or “phases”) during which a given fruit dominated the majority of the trials (Fig. 1B).

In order to maximize their gains, participants had to: (i) aggregate the evidence from each cue sequence to guess as quickly as possible the trial type; and (ii) use the structure of the task to infer, across trials, additional information about the probability of the outcome. Accordingly, we designed a computational model composed of two interacting modules: the first capturing the within-trial evidence accumulation and decision-making process (see *Decision model*) and the second tracking the trial-by-trial changes in trial types (see *Learning model*). Additionally, we implemented different hypotheses related to how the latter could interface with the former, *e*.*g*., by providing prior information or modulating the sensory integration dynamics (see *Learning-decision interactions*).

#### Decision model

This first module of the model describes how evidence is accumulated within a given trial, in order to reach and enact a decision about the basket type. Here, the model goal is to predict both the response (fruit type) and the CL as a function of the sequence of cues presented in the trial and the decision bias conferred by information from past trials.

Response is formalised as the ‘decision’ made by the observer after each cue presentation as a ternary choice between “cherries”, “watermelon” and “wait” (i.e., the choice to not yet respond, in order to observe additional cues and therefore accrue additional, decision-relevant information). Each trial is thus modelled as a set of (up to) 16 multinomial variables, with the corresponding observations represented as a vector: [*wait*^1^ *wait*^2^ … *wait*^*CL*−^ *choice NaN* …*NaN*].

At the core of the decision module is an accumulator that tracks the evidence in favour of the respective basket types (Eq. 1). Let us define *c* =[*c*^1^ *c*^2^ … *c*^*k*^ … *c*^16^]as the vector of cues composing a trial, e.g., with *c*^*k*^ = 0 if the *k*th cue was a cherries cue, and *c*^*ik*^ = 1 if it was a watermelon cue. We can then compute the accumulated log-evidence *LE*^*k*^ after each cue *k* as follows:

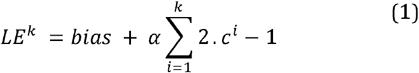

This function simply describes, e.g., counting positively (+1) the number of watermelon cues and negatively (-1) the cherries. Therefore, the value of the accumulator is increasingly positive when a majority of watermelon cues is being presented and, conversely, it becomes increasingly negative when more cherries are being presented. The *bias* term describes the potential prior influence of information from previous trials (see below). The accumulation rate *α* is fixed to 1 (but see *Learning-decision interactions*). From this, it straightforward to then compute the fruit cue-specific accumulated evidence after *k* cues:

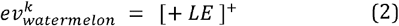

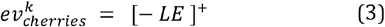

Where []^.+^ denotes the Heaviside step function, i.e., [*x*] ^.+^ = *x* if *x* > 0 and 0 otherwise.

While we could directly use this evidence to predict responses (e.g., by committing to choose when evidence goes over a fixed threshold), such a model could not account for effects such as the sense of urgency induced by the time-dependent reward function. Therefore, we introduce a ‘waiting’ signal (*w*) that counteracts the evidence accumulator:

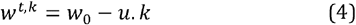

where *w*_0_ is the baseline tendency to wait and *u* a (positive) urgency signal that can speed up the need to respond as the trial approaches its end. As such, *w*_0_ and *u* are expected to be respectively negatively and positively related to impulsive responding.

Finally, we formalised the response as a categorical distribution with three possible events: {“wait”, “cherries”, “watermelon”}. The corresponding event probabilities (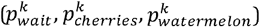) can then be expressed as a competition between the waiting signal *w*^*k*^ and the respective accumulators for the two cues, normalised to 1 with a softmax function:

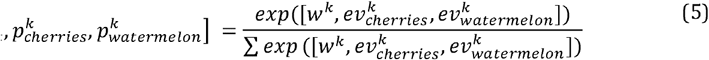

#### Learning model - Hierarchical Gaussian Filter (HGF)

As participants receive feedback directly after their response, they can infer what the actual trial type was. By observing and tracking those contingencies, they can build predictions about the most probable outcome at the next trial. We captured this learning process using the Hierarchical Gaussian Filter (HGF) model, a natural Bayesian extension of the reinforcement learning framework (Mathys et al., 2011, 2014; Iglesias et al., 2013; Diaconescu et al., 2014, 2017; Vossel et al., 2014; Lawson et al., 2017; Powers et al., 2017; Cole et al., 2020). For the sake of brevity, we will only outline the major components of the HGF and their notation and refer the reader to previous publications (e.g., Mathys, 2011) where it has been extensively described.

The belief at the first level of the learning model (*µ*_1_) corresponds to the observed environmental state or stimulus category (*x*_1_), in our case the type (dominant fruit) of a given trial. This belief is inferred on a trial-by-trial basis from the participant’s response (*y*) and the feedback they receive (*fbk*, equivalent to *x*_1_ in Mathys’ notation; see also the ‘Correct response’ blue dotted line in Fig. 1B):

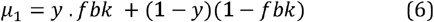

The HGF then implements a hierarchy of beliefs tracking the higher order statistics of the task structure: the second level (*x*_2_ ∼ *N* (*µ*_2_, *σ*_2_)) reflects the contingency expectation, that is the probability of the trial being of a certain type (*p* (*x*_1_)= *sig* (*µ*_2_)); the third and top level ((*x*_3_ ∼ *N* (*µ*_3_, *σ*_3_))tracks the (log-)volatility of the trial type, that is the uncertainty about the lower level beliefs due to the fluctuations of the trial type contingencies across the experiment. Practically, the volatility decreases when the trial probability (second-level belief) stays constant, as in the “stable” phases of the task.

Of note, fitting the HGF to each participant’s data allows recovery not only of their belief trajectories at the different levels, but also the (precision-weighted) prediction errors, which, at each trial, support the progressive updating of those beliefs. Those predictions errors, known to be encoded in specific neuromodulatory systems, can then be used in the analysis of the fMRI data to identify brain regions associated with the learning processes (see *fMRI analysis*).

#### Learning-decision interactions

The first type of interaction between the two modules relates to the use of predictions, learned from the task structure, as prior evidence for the decision phase. More precisely, as the HGF tracks the expected trial composition (*µ*_2_), it can provide a starting point for the trial-wise evidence accumulation process, prior to the presentation of any cues (Eq. 1)

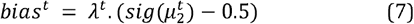

where *λ* is the weight of the learned expectation.

We also considered the possibility that the strength of this bias might be weighted by the uncertainty about the expected trial type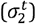 :

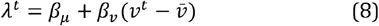

where *v*^*t*^ is the variance of the expected trial type (in the response space):

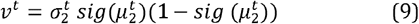

And 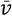 its grand average across the experiment, allowing orthogonalization of *β*_*µ*_ and *β*_*v*_.

We tested three hypotheses regarding the influence of the learned contingency belief on the trial-wise accumulation starting point: the null hypothesis is 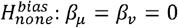, a first order influence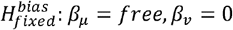, and a variance-weighted influence 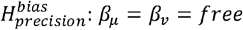.

In addition to this transfer of the predicted outcome belief to the decision module, we hypothesised that the accumulation process itself could be modulated by the volatility of the environment. For example, the general tendency to wait for more cues (*w*_0_) could increase with volatility (*µ*_3_) to ensure enough sampling in uncertain environment. Practically, we built four alternative variants (*H*^*volat*^) of the decision module where each of the parameters could be modulated by the volatility estimate:

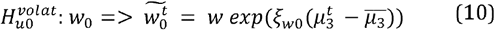

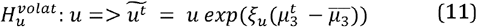

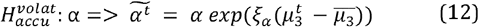

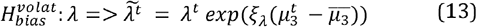

where “∼” denotes the volatility-modulated version of the parameter, replacing the corresponding fixed parameter in Eq. 1, 4, or 7, and 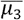is the grand average of *µ*_3_ across the experiment, ensuring orthogonality of the parameter influences. The null hypothesis 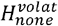 is that there is no influence of environment (*ξ*_*w*0_ = *ξ*_*u*_ = *ξ*_*α*_ = *ξ*_*λ*_ = 0).

#### Model space

The three hypotheses regarding the accumulator initial bias (*H*^*bias*^) can be factorially combined with the five hypotheses regarding the effects of volatility on evidence accumulation (*H*^*volat*^). However, as there is no possible volatility influence on bias if there is no bias at all (*λ* = 0), the combination of 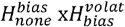 was excluded from the model space, which therefore contained 3 × 5 – 1 = 14 models (see Table 1, main text).

### Data analysis

#### Computational phenotyping

All the variants of our generative model of behaviour were implemented and fitted to each participant’s dataset using the Variational Bayes Analysis (VBA) toolbox (Daunizeau et al., 2014; https://mbb-team.github.io/VBA-toolbox/). The implementation of the HGF is identical to the one originally described by Mathys and colleagues (Mathys et al., 2011), which is provided as part of the TAPAS toolbox (Frässle et al., 2021). Priors are detailed Table S6.

To determine which combination of hypotheses best explains our sample behaviour, we then compared the resulting models *M*_1-14_ using Bayesian model selection (BMS) based on the free energy approximation of each model evidence (Stephan et al., 2009; Rigoux et al., 2014), as also implemented in the VBA-toolbox.

The maximum a posteriori estimates of the winning model were then collected from all participants for follow-up group-level analyses (see below). In addition, we extracted from this fitted model the trajectories of all the relevant beliefs and hidden states to be used as regressors in the fMRI analysis (see below), namely: (i) the accumulated log-evidence (*LE*^*t,k*^) and (ii) the waiting signal (*w*^*t,k*^), available for each cue of each trial, (iii) the initial bias of the accumulation *(λ* ^*t*^), (iv) the log-volatility estimate 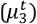, as well as (v-vi) the precision-weighted prediction errors at the second (*ε*_2_^*t*^) and third (*ε*_3_^*t*^) levels of the HGF, all computed for each trial.

### Behavioural analysis

In addition to the computational parameters described above, we derived purely data-driven metrics, to relate them to classical psychometric markers of impulsivity, and administered questionnaires, to assess their ecological validity.

#### Model-free metrics

As model-free markers of impulsivity, we first measured the choice latency (CL), calculated simply as the mean index (from 1 to 16) of the cue at which the participant responded. Shorter CL means the participant waited less and thus observed fewer fruits before committing to a choice, which could be interpreted through the lens of PRI. We also calculated the error rate, that is the percentage of trials in which the dominant fruit was not chosen. Again, higher error rates could also reflect suboptimal evidence accumulation and therefore some form of PRI.

Unfortunately, a speed-accuracy trade-off could create an interdependence between those measures, rendering them difficult to interpret independently. A well-established workaround is to calculate the so-called “I-score” and “E-score”. These composite scores combine error rates and reaction times (here CL) to provide theoretically untainted metrics of reflection impulsivity and cognitive efficiency, respectively (Salkind and Wright, 1977; Messer and Brodzinsky, 1981; Quednow et al., 2007b).

In order to test for effects of task structure on each of those measures, we also derived them independently for the volatile and stable phases of the task.

#### Questionnaires

We used several established questionnaires to assess impulsivity, including the BIS-11 questionnaire (Patton et al., 1995), which evaluates general impulsivity and its sub-factors (Nonplanning, Attentional, and Motor impulsiveness). We also administered the Monetary Choice Questionnaire (Kirby et al., 1999) to measure temporal delay discounting, and the UPPS Impulsive Behavior Scale (Whiteside and Lynam, 2001).

#### Analysis

When appropriate, we conducted Pearson bivariate correlation analyses to assess the interrelationships between these scales and psychometric measures and our computational measures of impulsivity. Next, to evaluate the influence of environmental uncertainty on impulsivity, we calculated, for the different metrics, the difference (Δ) between the volatile and stable phases of the task, using (one sample, two-sided) t-tests to establish significance, or Pearson’s correlation analyses to relate the volatility-induced behavioural changes with computational parameters or self-report scores.

### Image acquisition

Data were acquired on a 3 T Philips Achieva MRI scanner with an 8-channel head coil (Philips, Best, The Netherlands) at the Laboratory for Social and Neural Systems Research, University of Zurich. The quantity of whole-brain fMRI volumes acquired for each participant/session varied, as the concomitant behavioural paradigm was necessarily partially self-paced. T2*-weighted, sensitivity encoding (SENSE)-enabled echo-planar imaging (EPI) data sensitive to BOLD contrast were acquired during the task (TR = 2.66 s; TE = 36 ms; flip angle = 90°; 33 sequential ascending axial slices; in-plane resolution = 2.0 × 2.0 mm; slice thickness = 3.0 mm; slice gap = 0.6 mm; SENSE factor = 1.5; EPI factor = 63). Slice acquisition was angled upwards anteriorly by 20° from the axial plane. Magnetic equilibration was accounted for via scanner-automated dummy acquisition removal. High-resolution anatomical T1-weighted volumes were also acquired for each subject (TR = 8.3 ms; TE = 3.9 ms; flip angle = 8°; field-of-view 256 × 256 × 181; voxel size 1.0 × 1.0 × 1.0 mm; turbo field echo factor = 256).

For the behavioural task, stimuli were projected on a screen at the rear of the scanner that participants viewed through a mirror fitted to the top of the head coil. Participants’ cardiac and respiration rates were recorded during scanning with a 4-electrode electrocardiogram (ECG), pulse oximetry and a breathing belt. The partially self-paced nature of the task rendered it possible that task runs for some participants could extend longer in duration than many typical fMRI experimental runs. Therefore, for all participants, both task runs were split into two BOLD fMRI acquisitions, via the insertion of a ‘break’ at the same point in each run (between trials 133 and 134, thus just beyond halfway through the task, at the beginning of the second ‘stable’ pseudo-block). To minimise the disruption of natural learning processes, participants were informed that the task would continue from the same point at which it had stopped. Potential residual confounding effects of this break were controlled for in the fMRI analysis as described below (see *fMRI analysis*).

### fMRI data preprocessing

Preprocessing of fMRI data was performed using FSL 5.0 (Smith et al., 2004; https://fsl.fmrib.ox.ac.uk/). This incorporated the standard steps of slice-time correction, temporal high-pass filtering (128 seconds cut-off), realignment of individual volumes to correct for head motion, removal of non-brain tissue from the images and Gaussian smoothing at 5 mm full-width half-maximum (FWHM). Data from all fMRI sessions were transformed to a standard template space (Montreal Neurological Institute; MNI) for further analysis, firstly via boundary-based registration (Greve and Fischl, 2009) to the subject’s associated T1 anatomical volume, then via nonlinear registration to MNI standard space.

### fMRI analysis

Single-subject fMRI analyses were conducted using the general linear model (GLM) as implemented in SPM12 (Penny et al., 2007; http://www.fil.ion.ucl.ac.uk/spm/). Base regressors for the task were defined in terms of the onsets of each fruit cue stimulus in the decision period, the onset of the button press response for that trial (if applicable), and the onset of the outcome period, all of which were modelled as single events. In the case of the analysis incorporating the trial-wise bias weighting for prior information from the HGF model *(λ*_*t*_), additional events at the onset of the first cue stimulus and boxcar function covering the full decision period were also entered in the regression. The decision period base regressor was modelled along with two parametric modulator regressors encoding for the subject’s cue-wise waiting signal (*w*) and log-evidence accumulation which was squared (unsigned) to collapse across cue types (*LE*^2^) parameters (or one regressor encoding for the subject’s trial-wise bias weighting, *λ*_*t*_). Similar to previous work (Iglesias et al., 2013; Diaconescu et al., 2017; Cole et al., 2020) the outcome period regressor was loaded along with two parametric modulators encoding for the (absolute or signed) outcome-related precision-weighted prediction error (|*ε*_2_|, *ε*_2_) and the signed precision-weighted volatility-related prediction error (*ε*_3_) parameters (or three regressors in cases where *ε*_2_ was ‘split’ as described in *Supplementary Materials*). All parametric modulators were Fisher Z-normalised (zero mean, unit standard deviation) before entry into the GLM. Orthogonalization of regressors was not performed (this functionality was deactivated in SPM), as we did not want the results of our analyses to be impacted by the ordering of the regressors in the design matrix. No temporal or dispersion derivatives of any regressors were added to the subject-level GLMs. We performed physiological noise correction, based on RETROICOR (Glover et al., 2000) of ECG and respiratory measurements, using the physIO toolbox in TAPAS (Kasper et al., 2017) to compute 18 additional regressors that were included, along with six realignment parameters calculated during data preprocessing and representing head motion, as confound regressors in the GLM for each participant. Additional regressors generated by the FSL *fsl_motion_outliers* function flagged volumes with excessive movement (intensity spikes), and a last regressor encoded the parts before (0) and after (1) the break.

Group-level fMRI analyses were conducted, firstly using one-sample *t*-tests to examine overall neural encoding of computational PRI and other learning parameters, in terms of both positive and negative associative representations. Whole-brain results were thresholded using cluster-level family-wise error correction (p <.05, FWE), with a cluster-defining threshold of p <.001 (uncorrected). These analyses included contrast image data from both Gain and Loss task sessions, controlling for participant age, sex and mean attentional catch trial RT, along with a covariate representing the mean inter-trial interval. As an additional control to further minimise any effects of non-neuronal physiological noise on the results, analyses were conducted within a group mask consisting of a mean image of individual T1-weighted anatomical images nonlinearly registered to MNI space, then binarized and eroded using the *fslmaths* command in FSL. Secondly, we examined for differences in neural encoding of computational parameters between Gain and Loss task contexts using within-subject paired *t*-tests. Finally, due to our *a priori* interest in establishing links in humans between PRI behaviour and specific neurochemical systems, any results from the above contrasts occurring in midbrain regions of the brainstem were examined in terms of their overlap with masks of known dopaminergic, serotonergic and noradrenergic nuclei from independent studies (see Results).

### Psychophysiological interaction (PPI) analysis

In order to further address the neural mechanisms of PRI behaviour and their modulation by environmental volatility, we conducted a psychophysiological interaction (PPI) analysis (Friston et al., 1997; O’Reilly et al., 2012) using the preprocessed session-level fMRI data and confound regressors as described above in the initial model-based analysis. Here, instead of computational regressors of interest, the GLM included: (i) a boxcar regressor coded, respectively, as ‘-1’ or ‘1’ to describe whether the task was in a stable or volatile ‘pseudo-block’ (i.e., a blue or red period as depicted in Fig, 1B; the ‘psychological’ regressor), (ii) the mean time series of BOLD fMRI activation from a 5 mm radius sphere seed region surrounding the peak of the strongest cluster of group-level activation to the behavioural expression of state-level PRI (‘physiological’ regressor) and (iii) a regressor calculated as the multiplicative interaction between these first two (the PPI regressor). Statistical analysis was conducted using the FSL FEAT tool, with prewhitening switched on, orthogonalization of regressors switched off and no temporal derivatives considered. Group analysis of the contrast maps resulting from the subject-level PPI regressor analyses were conducted using one-sample and paired *t*-tests and ANCOVAs in SPM, in the same manner and with the same covariates as described in the initial group analysis above.

## Acknowledgments

We are grateful for support by the University of Zurich (UZH) and the UZH Clinical Research Priority Program (CRPP) “Molecular Imaging” (DMC, KES; UZH *Forschungskredit* K-82011-03-01, DMC), and the René and Susanne Braginsky Foundation (KES). We also acknowledge the valued study support and advice of our colleagues Karl Treiber, Thomas Baumgartner, Cornelia Schnyder, Eduardo Aponte, Aidan Makwana, Sara Tomiello, Sandra Iglesias, Helene Haker, Quentin Huys, and Klaas Enno Stephan.

