## Supplementary Materials for "A computational neuroimaging account of impulsive premature decision-making"

Table S1. Brain regions displaying significant BOLD activation changes in response to precision-weighted prediction errors during processing of decision outcomes. The upper panel displays regions that are significantly negatively activated in association with $|\varepsilon_{2}|$, while notably the exact same regions are also *positively* activated by $\varepsilon_{3}$ (any information that differs from that of the corresponding $|\varepsilon_{2}|$ cluster is written in **bold**), either significantly, or at trend levels^†^ (p < 0.10). The lower panel, conversely, displays the details of a single bilateral cluster positively activated in correlation with the signed version of $|\varepsilon_{2}|$ ($\varepsilon_{2}$). L = left hemisphere; R = right hemisphere.

| **Computational parameter (PE)** | **Brain region** | **Cluster size**  **(2 mm^3^ voxels)** | **P (cluster-level**  **FWE-corrected)** | ***t*-value of peak**  **voxel** | **x, y, z co-ordinates**  **(MNI space)** |
| --- | --- | --- | --- | --- | --- |
| $\vert\varepsilon_{2}\vert$, negative; and  $\boldsymbol{\varepsilon}_{\boldsymbol{3}}$**, positive** | L occipital fusiform gyrus | 81; **84** | 0.048; **0.046** | 5.81; **5.69** | -24, -82, -8 |
|  | L temporo-occipital fusiform cortex | 263; **206** | < 0.001 | 5.55; **5.61** | -30, -46, -18 |
|  | R temporo-occipital fusiform cortex^†^ | 113; **82** | 0.011; **0.051** | 5.53; **5.38** | 32, -48, -14; **32, -50, -16** |
|  | R occipital fusiform gyrus | 269; **371** | < 0.001 | 4.94; **5.24** | 30, -70, -10 |
|  | L superior lateral occipital cortex^†^ | 87; **80** | 0.036; **0.056** | 4.75; **4.92** | -26, -78, 20; **-28, -76, 20** |
|  | R superior lateral occipital cortex | 92; **99** | 0.029; **0.024** | 4.61; **4.72** | 28, -80, 18; **30, -80, 18** |
| $\varepsilon_{2}$, positive | Bilateral caudate | 227 | < 0.001 | 4.82 | -8 10, 8 |

Table S2. Brain regions with activation significantly correlated with only the *negative* part of the trajectory encoding signed outcome-related PEs ($\varepsilon_{2}$). STS = superior temporal sulcus; L = left hemisphere; R = right hemisphere.

| **Brain region** | **Cluster size**  **(2 mm^3^ voxels)** | **p (whole brain**  **FWE-corrected)** | ***t*-value of peak**  **voxel** | **x, y, z co-ordinates**  **(MNI space)** |
| --- | --- | --- | --- | --- |
| L occipital fusiform gyrus | 485 | < 0.001 | 5.92 | -22, -82, -8 |
| L superior lateral occipital cortex | 109 | 0.011 | 5.28 | -26, -78, 20 |
| R occipital fusiform gyrus | 221 | < 0.001 | 4.99 | 30, -78, -10 |
| R occipito-temporal cortex/STS | 116 | 0.008 | 4.91 | 56, -42, 0 |

Table S3. Brain regions significantly negatively encoding the weighting of bias towards prior (across-trial) information computed using the HGF ($\tilde{\lambda^{t}}$). AI = anterior insula; dlPFC = dorsolateral prefrontal cortex; IFG = inferior frontal gyrus; L = left hemisphere; R = right hemisphere.

| **Brain region** | **Cluster size**  **(2 mm^3^ voxels)** | **p (whole brain**  **FWE-corrected)** | ***t*-value of peak**  **voxel** | **x, y, z co-ordinates**  **(MNI space)** |
| --- | --- | --- | --- | --- |
| R occipital fusiform/lingual gyri | 735 | < 0.001 | 5.45 | 24, -78, -12 |
| L dorsal cerebellum | 90 | 0.026 | 4.94 | -24, -62, -32 |
| L superior occipital/occipito-parietal cortex | 273 | < 0.001 | 4.94 | -24, -80, 36 |
| L superior frontal cortex | 82 | 0.038 | 4.92 | -38, -10, 62 |
| L superior parietal lobule extending to inferior parietal cortex | 256 | < 0.001 | 4.82 | -34, -52, 52 |
| R occipito-temporal cortex | 259 | < 0.001 | 4.80 | 44, -70, 0 |
| R dlPFC | 112 | 0.009 | 4.75 | 42, 0, 32 |
| L occipital fusiform/lingual gyri ext. occipitotemporal cortex | 414 | < 0.001 | 4.74 | -20, -68, -6 |
| R IFG/AI | 166 | 0.001 | 4.69 | 46, 18, 4 |
| R superior occipital/occipito-parietal cortex | 78 | 0.046 | 4.63 | 28, -60, 54 |
| R superior lateral occipital cortex | 189 | < 0.001 | 4.52 | 32, -72, 28 |
| R superior parietal lobule ext. inferior parietal cortex | 103 | 0.014 | 4.35 | 28, -46, 52 |

Table S4. Brain regions with neurophysiological (BOLD fMRI) signal displaying functional connectivity with the right anterior insula (peak activation for the parametric modulator encoding impulsive decision-making behaviour, 1/$w$) that was significantly modulated by the task volatility (increased during volatile periods). dlPFC = dorsolateral prefrontal cortex; dmPFC = dorsomedial prefrontal cortex; OFC = orbitofrontal cortex; PCC = posterior cingulate cortex; vmPFC = ventromedial prefrontal cortex; L = left hemisphere; R = right hemisphere.

| **Brain region** | **Cluster size**  **(2 mm^3^ voxels)** | **p (whole brain**  **FWE-corrected)** | ***t*-value of peak**  **voxel** | **x, y, z co-ordinates**  **(MNI space)** |
| --- | --- | --- | --- | --- |
| Bilateral antero-medial cerebellum | 365 | < 0.001 | 6.52 | 6, -42, -28 |
| R dlPFC | 595 | < 0.001 | 6.25 | 28, 30, 50 |
| R temporopolar cortex extending to amygdala | 682 | < 0.001 | 6.05 | 44, 12, -32 |
| Bilateral PCC/precuneus, ext. precentral/postcentral gyri | 2538 | < 0.001 | 5.76 | 6, -38, 30 |
| Bilateral dmPFC/vmPFC ext. medial OFC | 1745 | < 0.001 | 5.68 | 0, 66, 26 |
| L opercular cortex ext. precentral gyrus | 600 | < 0.001 | 5.59 | -44, -14, 24 |
| R opercular cortex ext. precentral gyrus and posterior insula | 819 | < 0.001 | 5.57 | 50, -4, 18 |
| L occipital cortex/cuneus | 259 | < 0.001 | 5.43 | -2, -92, 28 |
| L temporopolar cortex ext. amygdala | 569 | < 0.001 | 5.25 | -48, 18, -26 |
| L precuneus/lingual gyrus | 105 | 0.004 | 5.25 | -16, -48, 6 |
| R mid-cingulate cortex | 101 | 0.005 | 5.24 | 4, -8, 44 |
| R inferior parietal cortex/angular gyrus | 290 | < 0.001 | 5.22 | 60, -50, 40 |
| L lateral OFC | 73 | 0.030 | 4.92 | -38, 42, -10 |
| R caudate ext. white matter | 88 | 0.011 | 4.81 | 28, 2, 26 |
| R dlPFC | 221 | < 0.001 | 4.69 | 48, 22, 42 |
| L superior temporal gyrus | 91 | 0.009 | 4.51 | -56, -40, 12 |
| R lateral occipital/inferior parietal cortex | 87 | 0.012 | 4.44 | 28, -60, 40 |
| R lateral OFC | 71 | 0.035 | 4.30 | 34, 42, -12 |

Table S5**.** Model-based PRI, HGF and other non-model-based learning parameter descriptive statistics (mean ± s.d.) calculated separately and compared for Gain and Loss task contexts (n = 23). Accumulation rate $\alpha$ is not reported on here as a MAP estimate, as it has a fixed prior variance (see Table S1). However, $\alpha$ informs a dynamic accumulation rate variable ($\tilde{\alpha^{t}}$) weighted by trial-wise volatility in the winning model (here this weighting parameter is reported on and labelled as $\xi_{\alpha}$). CL = choice latency (mean number of stimuli waited across the task). NB: due to the normalisation procedure (Fisher Z-scoring) used in the calculation of these parameters, mean I- and E-scores across participants for both the Gain and Loss task conditions were zeroed and thus it is not possible to compare them statistically across task.

|  | Gain | Loss | *t* | p |
| --- | --- | --- | --- | --- |
| $w_{0}$ | 4.65 ± 1.47 | 4.77 ± 1.55 | 0.57 | 0.58 |
| $u$ | 4.97 ± 3.37 | 5.79 ± 5.09 | 1.29 | 0.21 |
| $\lambda$ | 2.79 ± 1.79 | 3.13 ± 1.91 | 1.03 | 0.31 |
| $\xi_{\alpha}$ | 3.50 ± 4.45 | 4.45 ± 8.76 | 0.44 | 0.66 |
| $\kappa$ | 0.94 ± 0.37 | 0.99 ± 0.36 | 0.45 | 0.66 |
| $\theta$ | 0.09 ± 0.09 | 0.11 ± 0.09 | 0.59 | 0.57 |
| CL | 4.98 ± 1.59 | 5.00 ± 1.70 | 0.08 | 0.94 |
| I-score | 0 ± 1.82 | 0 ± 1.77 | - | - |
| E-score | 0 ± 0.83 | 0 ± 0.93 | - | - |

Table S6. Priors on model parameters.

| **Model part** | **Parameter** | **Prior mean** | **Prior variance** |
| --- | --- | --- | --- |
| Between-trial HGF (perceptual) module | κ | 0 | 0.8 |
|  | $\omega$ | -4 | 0 |
|  | 𝜗 | log(0.05) | 1 |
|  | $\mu_{2}^{(k=0)}$ | 0 | 0.2 |
|  | $\sigma_{2}^{(k=0)}$ | log(0.3) | 0.5 |
|  | $\mu_{3}^{(k=0)}$ | 0 | 0 |
|  | $\sigma_{3}^{(k=0)}$ | log(4) | 0 |
| Within-trial accumulator (decision) module | $\alpha$ | 1 | 0 |
|  | $u_{0}$ | log(5) | 1 |
|  | $u_{K}$ | log(2) | 1 |

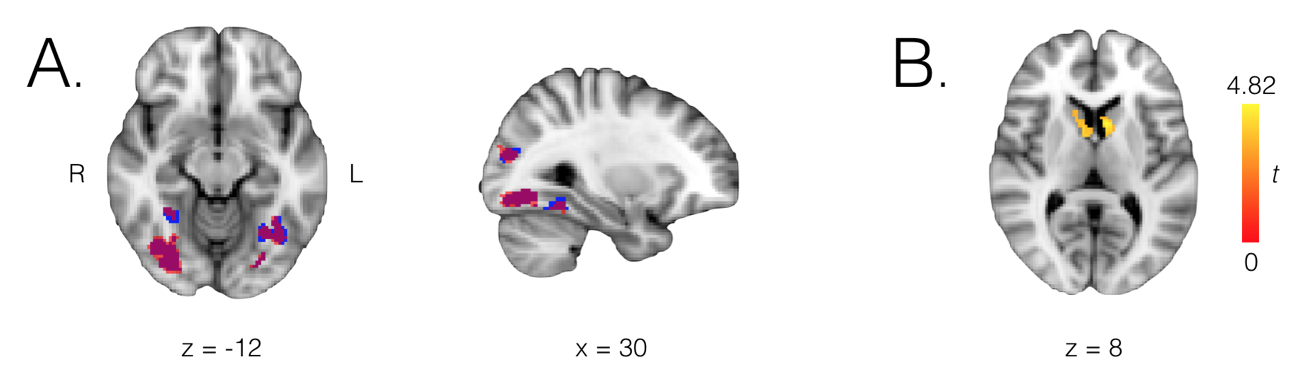

Figure S1. Brain regions activated in correlation with precision-weighted prediction error (PE) signals during monitoring of decision outcomes. (A) Regions significantly (whole-brain cluster-level FWE-corrected p < 0.05) negatively activated by outcome-related PEs ($|\varepsilon_{2}|$, blue) or significantly positively activated by volatility-related PEs ($\varepsilon_{3}$, red), overlaid as binary clusters with overlap displayed (purple). (B) Bilateral caudate regions display positive activation by an alternative *signed* version of the outcome-related PE (±$\varepsilon_{2}$); colour bar represents *t*-statistic effect sizes shown in brain image overlay. R = right, L = left. NB: axial slices are displayed in line with radiological orientation conventions (L/R hemisphere flipped in figure). Binary and statistical maps overlaid on background image of a T1 template in MNI152 standard space.

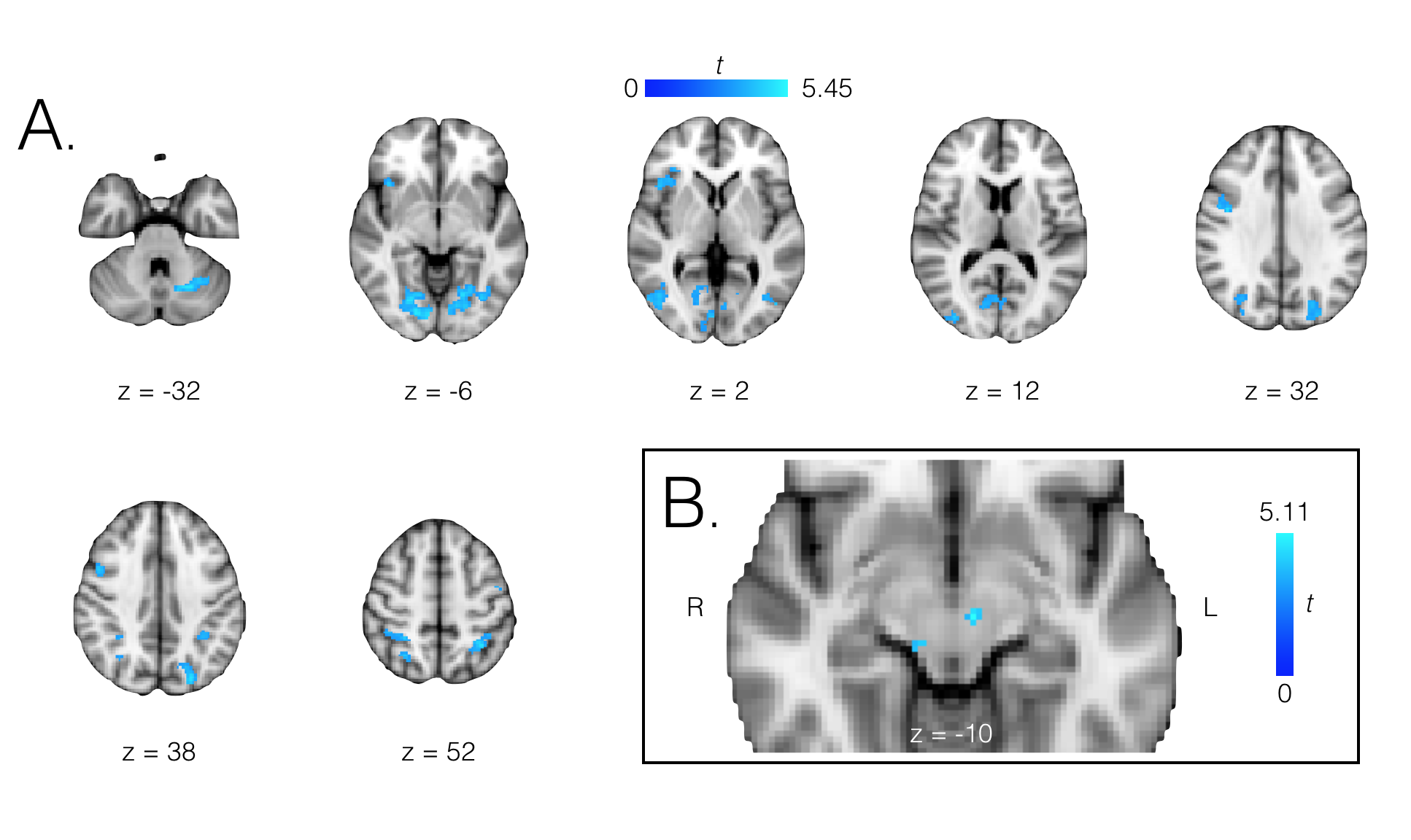

Figure S2. Brain regions with activation significantly negatively associated with the bias weighting of prior information during decision-making ($\lambda^{t}$). (A) Whole-brain cluster-level FWE-corrected p < 0.05, cluster-defining voxel-level threshold p < 0.001; and (B) small volume corrected (p < 0.05) for an *a priori* mask of the substantia nigra/ventral tegmental area (SN/VTA) of the dopaminergic midbrain. R = right, L = left. NB: axial slices are displayed in line with radiological orientation conventions (L/R hemisphere flipped in figure). Colour bars represent *t*-statistic effect sizes shown in brain image overlays. Statistical maps overlaid on background image of a T1 template in MNI152 standard space.

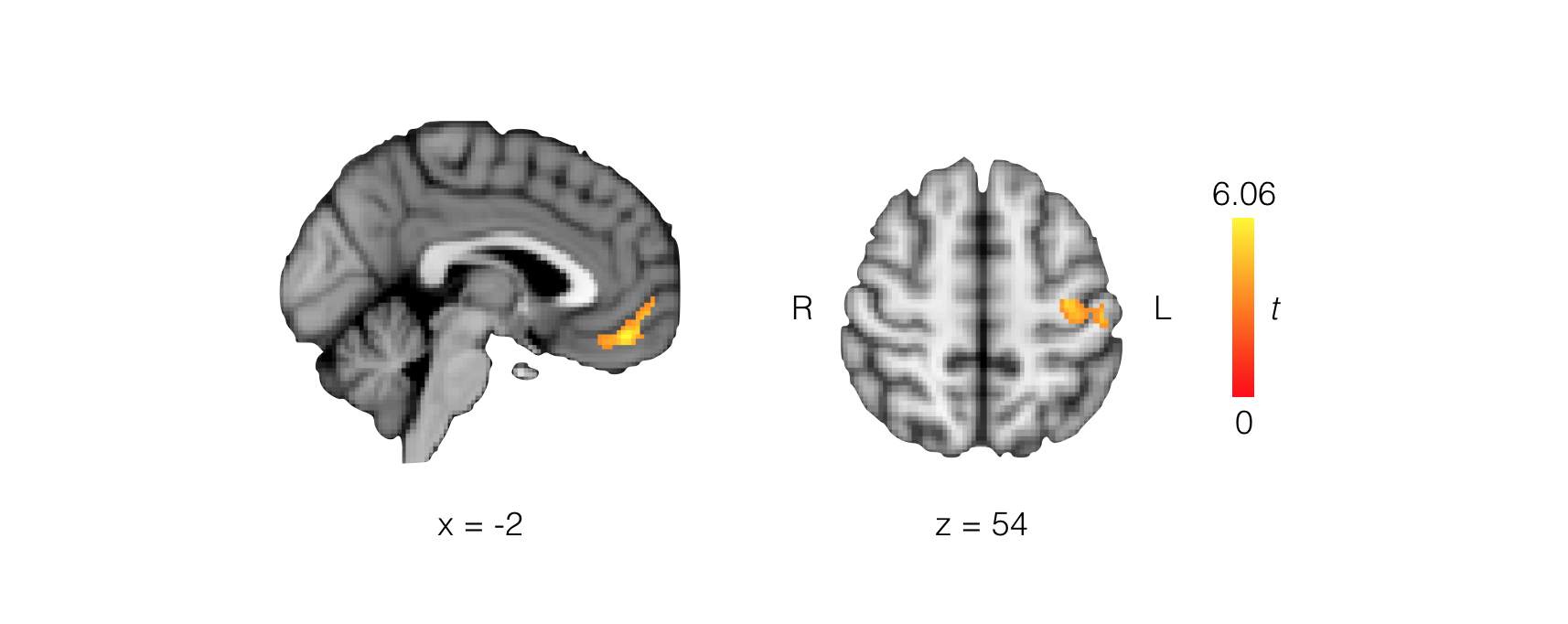

Figure S3. Brain regions with activation significantly positively associated with the precision weighting ($\sigma_{2}$) parameter alone, separately from the outcome-related PE, during monitoring of decision outcomes (whole-brain cluster-level FWE-corrected p < 0.05, cluster-defining voxel-level threshold p < 0.001). R = right, L = left. NB: axial slices are displayed in line with radiological orientation conventions (L/R hemisphere flipped in figure). Colour bar represents *t*-statistic effect sizes shown in brain image overlays. Statistical maps overlaid on background image of a T1 template in MNI152 standard space

**Results – supplementary**

*Model-based fMRI: precision-weighted prediction errors*

Initial one-sample *t*-test analyses examining outcome-related ($|\varepsilon_{2}|$, absolute values) and volatility-related ($\varepsilon_{3}$, signed values) PEs, pooled across Gain and Loss task contexts, revealed significant activations (whole-brain cluster-level FWE-corrected p < 0.05, cluster-defining voxel-level threshold p < 0.001) associated with both computational measures in regions of visual cortex (Fig. S2A; Table S4). Importantly, these activations were negatively correlated with $|\varepsilon_{2}|$ and positively correlated with $\varepsilon_{3}$. No significant effects were found in terms of positive associations with $|\varepsilon_{2}|$ or negative associations with $\varepsilon_{3}$. However, a repetition of this analysis with a *signed* outcome-related PE ($\varepsilon_{2}$) revealed a significant activation in bilateral caudate regions (Fig. S2B; Table S5), while no significant negative associations with $\varepsilon_{2}$ were found in any regions, including those of visual cortex that were deactivated by *absolute* outcome-related PEs ($|\varepsilon_{2}|$).

A second adaptation of this analysis, which involved testing the signed PE ($\delta_{1}$) and the precision-weighting ($\sigma_{2}$) as separate parametric modulators in the fMRI design (rather than combining these to form $|\varepsilon_{2}|$ or $\varepsilon_{2}$) found that the above result, showing $\varepsilon_{2}$ to be significantly represented in bilateral dorsal caudate, was driven by the neural encoding of the PE itself ($\delta_{1}$), although the cluster and effect sizes (*t* = 4.57; p = 0.001; x = -8, y = 10, z = 8; 176 voxels) were somewhat less strong than those of the combined $\varepsilon_{2}$ measure (see Table S6). The $\sigma_{2}$ precision weighting measure alone, however, exhibited significant positive encoding in regions of bilateral ventromedial prefrontal/medial orbitofrontal cortex (*t* = 6.06; p = 0.001; 0, 46, -16; 172 voxels) and left precentral gyrus (*t* = 5.12; p = 0.005; -34, -16, 54; 127 voxels; Fig. S3). A third and final adaptation of the outcome-related precision-weighted PE analysis, where the signed PE ($\varepsilon_{2}$) was split into two parametric modulator variables encompassing either its positive or its negative parts (by fixing at zero any sections of the trajectory that were, respectively, below or above zero), conversely did not reveal any significant activation of the caudate by either part. In fact, this analysis revealed only that, when considered in isolation, the negative part of the signed PE correlated significantly with activations in visual cortex regions (Table S6), which were largely comparable to the aforementioned regions negatively activated by the *absolute* outcome-related PE, $|\varepsilon_{2}|$ (and positively activated in line with the signed volatility-related PE, $\varepsilon_{3}$; see also Table S8). Together, these results show that the complete, signed outcome-related prediction error, $\varepsilon_{2}$, is the signal necessary for evoking outcome-related, PE-modulated activations in bilateral caudate regions during the task. Similarly to the analysis of computational PRI parameters, no differences in the neural representations of any precision-weighted PE parameters were found between Gain and Loss task contexts, neither were any significant main effects of sex or task contest × sex interactions identified.

*Model validation*

**Parameter recovery**

In order to assess the reliability of our parameter estimation procedure, we first simulated 100 datasets from the winning model using random parameters uniformly sampled from a range encompassing the values estimated from the participants’ behaviour. We then applied our model fitting routine to each synthetic dataset and collected the parameter estimates. Finally, for each parameter, we plotted and quantified (Pearson r) the correlation between the simulated (ground truth) and the estimated value. As shown Figure S4, all parameters can be reliably recovered.

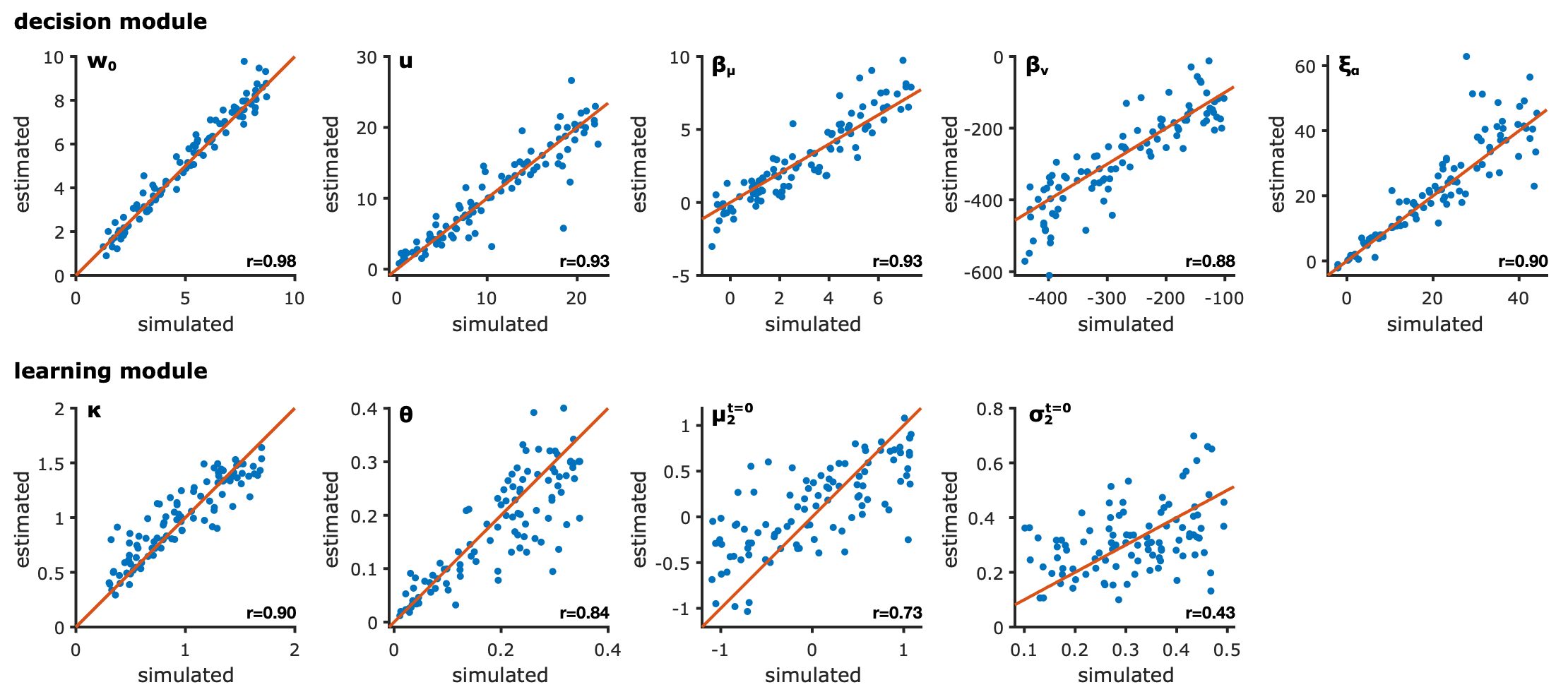

***Figure S4: parameter recovery.*** *Correlation between simulated and estimated parameter values for the winning model. Statistics: Pearson’s r.*

*Model recovery*

Similarly, we ensured that our statistical pipeline could correctly discriminate our different hypotheses. To this end, we first simulated 100 datasets from each of the 14 models of our model space. For each synthetic dataset, we then computed the model evidence for all the competing models and identified the winning one. Finally, we calculated for each simulated model the proportion of model simulations being identified correctly or not, forming the so-called confusion matrix (figure S5). While some models were misclassified, those errors were primarily related to the H^bias^ hypotheses, with “precision-weighted” models being identified as simpler “fixed” models, and did not relate to the H^volat^ hypotheses, i.e., the identification of the effect of volatility on decision parameters.

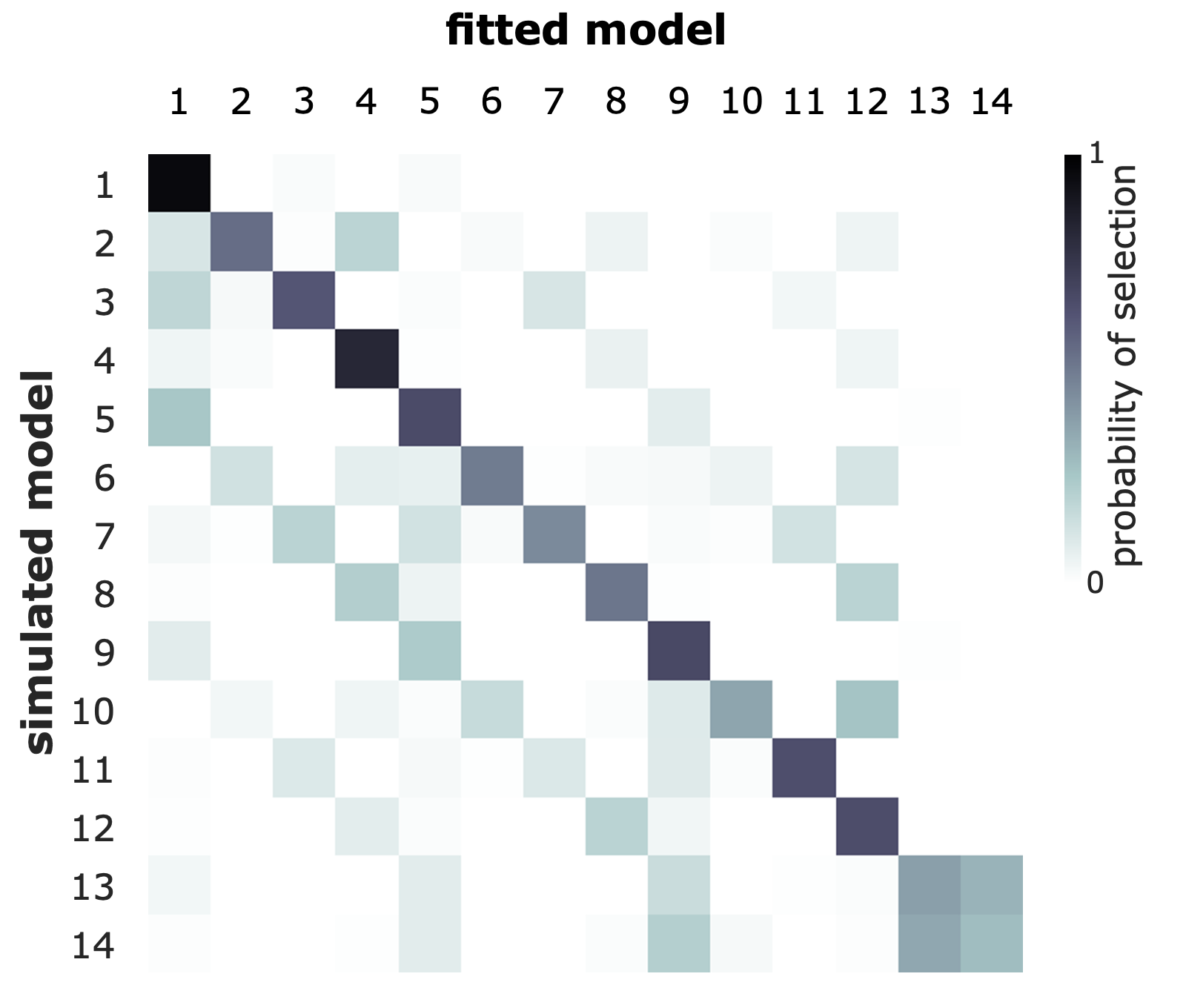

***Figure S5: model recovery.*** *Confusion matrix showing in what proportion (shade) datasets simulated from the respective hypotheses (lines) are attributed to the different models (column). The diagonal represents correct attributions.*

**References.**

Baker SC, Konova AB, Daw ND, Horga G (2019) A distinct inferential mechanism for delusions in schizophrenia. Brain 142:1797–1812.

Behrens TEJ, Woolrich MW, Walton ME, Rushworth MFS (2007) Learning the value of information in an uncertain world. Nat Neurosci 10:1214–1221.

Cole DM, Diaconescu AO, Pfeiffer UJ, Brodersen KH, Mathys CD, Julkowski D, Ruhrmann S, Schilbach L, Tittgemeyer M, Vogeley K, Stephan KE (2020) Atypical processing of uncertainty in individuals at risk for psychosis. NeuroImage Clin 26:102239.

Diaconescu AO, Mathys C, Weber LAE, Kasper L, Mauer J, Stephan KE (2017) Hierarchical prediction errors in midbrain and septum during social learning. Soc Cogn Affect Neurosci 12:618–634.

Iglesias S, Mathys C, Brodersen KH, Kasper L, Piccirelli M, den Ouden HEM, Stephan KE (2013) Hierarchical prediction errors in midbrain and basal forebrain during sensory learning. Neuron 80:519–530.

Kirby KN, Petry NM, Bickel WK (1999) Heroin addicts have higher discount rates for delayed rewards than non-drug-using controls. J Exp Psychol Gen 128:78–87.

Massar SAA, Lim J, Sasmita K, Chee MWL (2016) Rewards boost sustained attention through higher effort: A value-based decision making approach. Biol Psychol 120:21–27.

Mathys C, Daunizeau J, Friston KJ, Stephan KE (2011) A bayesian foundation for individual learning under uncertainty. Front Hum Neurosci 5:39.

Paliwal S, Petzschner FH, Schmitz AK, Tittgemeyer M, Stephan KE (2014) A model-based analysis of impulsivity using a slot-machine gambling paradigm. Front Hum Neurosci 8:428.

Patton JH, Stanford MS, Barratt ES (1995) Factor structure of the barratt impulsiveness scale. J Clin Psychol 51:768–774.

Rothenhoefer KM, Hong T, Alikaya A, Stauffer WR (2019) Rare Rewards Amplify Dopamine Learning Responses. bioRxiv.

Trueblood JS, Busemeyer JR (2011) A Quantum Probability Account of Order Effects in Inference. Cogn Sci 35:1518–1552.

Turner DC, Aitken MRF, Shanks DR, Sahakian BJ, Robbins TW, Schwarzbauer C, Fletcher PC (2004) The role of the lateral frontal cortex in causal associative learning: exploring preventative and super-learning. Cereb Cortex 14:872–880.
